# Vitamin D differentially modulates effector and regulatory T-cell migration across the blood-brain barrier

**DOI:** 10.64898/2025.12.09.692504

**Authors:** Sasha Soldati, Alexander Bär, Felix Blumer, Arindam Pal, Saniya Kari, Mykhailo Vladymyrov, Ahmad Danial, Hideaki Nishihara, Justin Killick, Adrian Mossu, Elisa Bouillet, Sara Barcos, Manon Galoppin, Maria Rosito, Urban Deutsch, Federica Sallusto, Mireia Sospera, Roland Martin, Fabien Gosselet, James L. McGrath, Eric Thouvenot, Anne L Astier, Britta Engelhardt

**Affiliations:** Theodor Kocher Institute, University of Bern, Bern, Switzerland; Univ Toulouse, Toulouse Institute for Infectious and Inflammatory Diseases (Infinity), INSERM UMR1291 - CNRS UMR5051, 31024 Toulouse cedex 3, France; Department of Biomedical Engineering, University of Rochester, Rochester, NY, USA; Centre for Inflammation Research, University of Edinburgh, Edinburgh, UK; Department of Neurology, Nîmes University Hospital, University Montpellier, Nîmes, France; Institute of Microbiology, ETH Zürich, Zürich, Switzerland; Institute for Research in Biomedicine, Università della Svizzera italiana, Bellinzona, Switzerland; Institute for Experimental Immunology, University of Zürich, Zürich, Switzerland; Blood-Brain Barrier Laboratory, University of Artois, Lens, France

**Author notes:** **correspondence:** Dr. Britta Engelhardt, Theodor Kocher Institute, University of Bern, CH - 3012 Bern, Switzerland, Dr. Anne L. Astier Infinity, INSERM U1291, CNRS U5051 Université de Toulouse 31024 Toulouse cedex France. Present address: University of Texas at Dallas, Department of Bioengineering, and University of Texas Southwestern Medical Centre, Department of Biomedical Engineering, Dallas TX, USA. Present address: Institut Cochin, Inserm U1016 - CNRS UMR8104 - Université Paris Cité, Paris. Present address: Yamaguchi University, Department of Neurology and Clinical Neuroscience, Yamaguchi, Japan (Hideaki Nishihara). Present address:Link Campus University, Department of Life Sciences, Health and Health Professions. Rome, Italy. Equal contribution (Anne L Astier, Britta Engelhardt). These authors contributed equally to this work.

**Keywords:** Vitamin D, multiple sclerosis, T cell migration, blood-brain barrier, regulatory T cells

## Abstract

Multiple sclerosis (MS) is an inflammatory disease of the CNS influenced by a combination of genetic predisposition and environmental factors. Vitamin D (VitD) deficiency is considered a major risk factor for MS. While VitD is associated with immunomodulatory roles, the exact mechanisms by which VitD protects from disease are still largely unknown. CD4 T cells play a key role in MS pathogenesis with autoimmune effector T cells (Teff) infiltrating the CNS across the blood-brain barrier (BBB) and regulatory T cells (Treg) displaying impaired functions.

Here we show that treatment of human CD4 T cells with the active form of VitD (1,25-Dihydroxyvitamin D3; 1,25(OH)2D3) decreased cell-surface expression of α4β1- and αLβ2-integrins on Teff but not Treg and reduced Teff adhesion to their endothelial ligands VCAM-1 and ICAM-1. By employing live cell imaging, we observed that VitD treatment reduced arrest of Teff but not Treg to the BBB as well as ICAM-1 and VCAM-1 under physiological flow *in vitro* and differentially affected post-arrest behaviour of Teff versus Treg on the BBB under physiological flow. Furthermore, VitD treatment favoured the migration of Treg over Teff across the BBB under static and flow conditions *in vitro. In vivo* live cell imaging showed that VitD reduced T cell arrest on the inflamed BBB during autoimmune neuroinflammation. Finally, VitD also reduced expression of integrins mediating CNS homing on pathogenic CD4 T cells isolated from the CSF of persons with MS (PwMS).

As VitD treatment did not alter barrier properties or adhesion molecule profile of our BBB model we propose a beneficial effect of VitD supplementation in PwMS by reducing CNS trafficking of pro-inflammatory T cells while leaving CNS entry of Treg unaffected.

## Introduction

Multiple sclerosis (MS) is a chronic inflammatory disease of the central nervous system (CNS)^1,2^ with genetic predisposition and environmental triggers.^3–5^ Among the latter, low sun exposure and vitamin D (VitD) deficiency have been shown as major risk factors for the development of many autoimmune diseases, including MS.^6^

It is well established that low 25-hydroxyvitamin D3 (25(OH)D3; calcidiol) serum levels correlate with an increased risk of developing MS. Persons with MS (PwMS) have lower levels of circulating VitD than healthy controls^7,8^ and their VitD serum levels inversely correlate with disease activity and disability progression suggesting a protective role of VitD in MS.^9,10^ Indeed, the recent randomized double-blind placebo-controlled trial (D-Lay-MS) aiming at supplementing patients with high dose of VitD as a monotherapy at their first clinical symptoms (trial NCT01817166) showed that VitD significantly reduced disease activity in early MS^11^, strongly supporting the hypothesis that VitD contributes to ameliorating MS pathogenesis. The beneficial effect of VitD in PwMS could result from its known neuroprotective and immunomodulatory functions^12^, as the vitamin D receptor (VDR) is expressed in CNS-resident cells and different immune cell subsets^13,14^, including activated T cells.^15^ VitD exerts its immunomodulatory role in part by reducing inflammatory cytokines^16^ while enhancing IL-10.^17^ Furthermore, VitD inhibits differentiation of Teff while promoting differentiation of conventional CD25^+^Foxp3^+^ regulatory T cells (Treg)^18^ and type I regulatory T cells (Tr1)^17^, suggesting an overall anti-inflammatory effect of VitD.

One of the pathological hallmarks and precondition of MS is immune cell infiltration into the CNS leading to neuroinflammation and demyelination.^19^ Immune cell migration into the CNS has been extensively studied in experimental autoimmune encephalomyelitis (EAE), an animal model of autoimmune neuroinflammation.^20,21^ T cell migration across the BBB is a multi-step process that involves the sequential interaction of different adhesion- and signaling molecules on the BBB and T cells.^22^ After initial capture and rolling, mediated by P-selectin glycoprotein ligand (PSGL)-1, T cells arrest on the BBB which is mediated by α4β1- and αLβ2-integrins (also known as VLA-4 (CD49d/CD29) and LFA-1 (CD11a/CD18), respectively), binding to vascular cell adhesion molecule (VCAM)-1 and intercellular adhesion molecule (ICAM)-1, their respective endothelial ligands. After polarizing and crawling against the direction of blood flow on endothelial ICAM-1 and ICAM-2, T cells cross the BBB either paracellularly, between three or two adjacent endothelial cells, or transcellularly in a pore through the endothelial cell-body.^23^ Importantly, therapeutic targeting of immune cell migration across BBB, with natalizumab (NTZ), a humanized function-blocking anti-α4-integrin antibody, efficiently blocks disease activity and new MRI lesions in relapsing remitting (RR)MS.^24^ However, a serious and not rarely fatal side effect, progressive multifocal leukoencephalopathy (PML), has been observed. This experience raises significant concerns about the safety of therapies targeting T-cell trafficking as these might also affect the migration of beneficial T-cell subsets maintaining immune surveillance of the CNS. It is therefore key to clearly determine what affects migration of effector and/or regulatory T cells and to identify new therapeutic targets to specifically inhibit the migration of pathogenic T cells into the CNS.

VitD suppresses the development and progression of EAE^25^ by modulating effector cytokines and decreasing the infiltration of T cells into the CNS.^26,27^ Furthermore, active VitD modulates adhesion molecules on activated human CD4 T cells which could affect their trafficking properties.^28^ However, the impact of VitD on the migration of the distinct human T cell subsets involved in MS across the BBB has not been investigated. By employing *in vitro* and *in vivo* live cell imaging, we investigated the effect of 1,25(OH)2D3 on the multi-step migration of both effector and regulatory T-cell subsets across the BBB in the context of neuroinflammation. Our study provides direct evidence of an effect of VitD on T-cell migration across the BBB by reducing the cell-surface expression of α4β1- and αLβ2-integrins on CD4 T cells and their adhesion to VCAM-1 and ICAM-1 *in vitro* and reducing arrest of CD4 T cells on the inflamed BBB endothelium under physiological flow conditions both *in vitro* and *in vivo*. Furthermore, we found a differential effect of VitD on the migration of Teff and Treg, favoring the migration of Treg over Teff across the BBB *in vitro*. Our study provides novel insights into the immunomodulatory role of VitD and supports the notion that VitD supplementation could contribute to better clinical outcomes in PwMS by reducing the infiltration of Teff but not Treg into the CNS.

## Materials and methods

### Mice

Mice were housed in individually ventilated cages under specific pathogen-free conditions at 22°C with free access to chow and water. All animal procedures were approved by the Veterinary Office of the Canton Bern (permit no. BE33/20 and BE98/20) and are in line with institutional and standard protocols for the care and use of laboratory animals in Switzerland. VE-Cadherin-GFP knock-in C57BL/6J mice are described in^29^.

### Healthy donors and PwMS

Buffy coats from healthy individuals were purchased from the Swiss Red Cross (Interregionale Blutspende SRK, Bern, Switzerland; project number P_172) while CSF samples from PwMS were obtained under ethics protocol EC-No. 2014-0699 (approved by the Ethics Committee of the Canton of Zurich on 27th February 2015). Detailed information of the PwMS can be found in **Supplementary Table 1**.

### T-cell culture

PBMCs were isolated from buffy coats of healthy donors by Ficoll-Paque^TM^ Plus (Cytiva) density gradient and CD4 T cells purified by negative selection (EasySep™ STEMCELL technologies, #17952) following the provider’s instructions. CD4 T-cell purity analyzed by flow cytometry was always above 95%. The employed flow cytometry antibodies are listed in **Supplementary Table 2**. CD4 T cells were subsequently activated with anti-human CD3 (OKT-3, Biolegend #317326) and anti-human CD28 (CD28.2, Biolegend, #302902) for 5 days in T-cell culture medium (TCM) (RPMI-1640 (Gibco), 10% (v/v) heat inactivated fetal bovine serum (FBS, Hyclone), 2 mM L-Glutamine (Gibco), 1% (v/v) MEM Non-Essential Amino Acids Solution (Gibco), 1 mM Sodium Pyruvate (Gibco), and 0.05 mM β-Mercaptoethanol (Grogg Chemie AG), 10 IU/mL Penicillin-Streptomycin (Gibco), 100 μg/mL Kanamycin Sulfate (Gibco), 10 IU/mL recombinant human interleukin 2 (IL-2)) in the presence of 100 nM 1,25(OH)2D3 or same amount (v/v) of vehicle control (EtOH) as previously described.^28^

The different Th subsets (Th1, Th2, Th1*, and Th17) were isolated by flow cytometry, and expanded as previously described.^30^ Briefly, purified human CD4 T cells from buffy coats of healthy donors or from the CSF of MS patients were sorted by FACS according to their specific cell-surface expression of chemokine receptors (CCR6^−^CXCR3^+^CCR4^−^ for Th1; CCR6^+^CXCR3^+^CCR4^−^ for Th1*; CCR6^−^CXCR3^–^CCR4^+^ for Th2; CCR6^+^CXCR3^−^CCR4^+^ for Th17, purity > 90%, **Supplementary Table 3**). Th cells were subsequently expanded for 20 days with periodic re-stimulation with 1 μg/mL phytohemagglutinin, 500 IU/mL IL-2 and irradiated allogeneic PBMCs in TCM. Treg were purified using the CD4^+^CD127^low^CD25^+^ Treg Isolation kit (EasySep™, STEMCELL technologies) (purity > 80%, **Supplementary Table 4)** and expanded for 14 days using a Treg expansion kit (Myltenyi Biotec). The distinct Th subsets were cultured for 5 days in the presence of 100 nM 1,25(OH)2D3 or same amount (v/v) of vehicle control (EtOH). T cells were then labelled if needed with 1 μM CellTracker™ Green (CMFDA Dye, Life technologies) or Deep Red (Life technologies) for 30 min at 37°C (5% CO2) prior to the experiments.

### Brain-like endothelial cells as an *in vitro* model of the BBB

Brain-like endothelial cells (BLECs) were used as a human *in vitro* model of the BBB as previously described ^31,32^. The protocol for the handling of human tissues and cells was authorized by the French Ministry of Higher Education and Research (CODECOH Number DC2011-1321) and all parent’s infants gave their approval.

For permeability assays, trans-endothelial electrical resistance (TEER) measurement, immunostaining, and flow cytometry analysis, CD34^+^ endothelial cells were grown to confluency on Matrigel^TM^-coated filter inserts (PC membrane, pore size 0.4 μm; Costar, 3401) in co-culture with brain pericytes for 6 days. BLECs were then treated with 100 nM 1,25(OH)2D3 or same amount (v/v) of vehicle control (EtOH) for 2 additional days.

For *in vitro* live-cell imaging, CD34^+^ endothelial cells were cultured to confluency on Matrigel^TM^-coated nanoporous silicon nitride (NPN) membranes of μSiM-CVB flow chambers in co-culture with bovine pericyte-conditioned medium for 6 days.

BLECs were stimulated or not with 76 IU/mL of recombinant hTNFα (R&D systems, 210TA) and 20 IU/mL recombinant hIFNγ (R&D systems, 285IF) for 16-24 h at 37°C (5% CO2) prior experiments.

### Cell-surface adhesion molecule expression analysis by flow cytometry

BLECs were cultured to confluency as described above. The cell-surface molecule expression of VE-cadherin, PECAM-1, VCAM-1, ICAM-1, ICAM-2 and CD99 was analyzed by flow cytometry as previously described^31^. Antibodies are listed in **Supplementary Table 5**. Human CD4 T cells, CD4 Th cells (Th1, Th2, Th1*, and Th17) and Treg were cultured and treated as described above. The cell-surface molecule expression of α4-, α5-, α6-, αL-, β1, β2- and β7-integrins, PSGL-1, PECAM-1, ALCAM, CD99 and L-selectin was analyzed by flow cytometry (antibodies listed in **Supplementary Table 6)**.

### Immunofluorescence staining of BLECs

BLECs were cultured to confluency on filter inserts as described above. Immunofluorescence staining for VCAM-1, ICAM-1, ZO-1, VE-cadherin, claudin-5, and fibronectin on BLECs was performed as previously described.^31^ In brief, primary antibodies for ICAM-1 and VCAM-1 were added to live cells and incubated for 15 min at 37°C (5% CO2). After washing, cells were fixed with 1% (w/v) paraformaldehyde and blocked with 5% (w/v) skimmed milk in PBS for VDR, ZO-1, VE-cadherin, and fibronectin staining. For claudin-5 staining cells were fixed with ice cold methanol and blocked with blocking buffer (5% skimmed milk, 0.3% Triton-X, 0.04% sodium azide in Tris-buffered saline (TBS) pH 7.4). BLECs were then incubated with primary antibodies in blocking buffer for 1 hour at RT. After washing with PBS, BLECs were incubated with fluorochrome-labelled secondary antibodies in blocking buffer for 1 hour at RT. Nuclei were stained with 1 μg/mL DAPI. After washing with PBS, filters inserts were mounted with Mowiol (Sigma-Aldrich) to a glass slide and imaged using a Nikon Eclipse E600 microscope connected to a Nikon Digital Camera DXM1200F with Nikon NIS-Elements BR3.10 software (Nikon, Egg, Switzerland). Antibodies used are listed in **Supplementary Table 7**.

### Assessment of blood-brain barrier integrity *in vitro*

BLECs were cultured to confluency on filter inserts as described above. *In vitro* permeability of BLECs monolayers to Lucifer Yellow (LY; 457 Da) (Sigma-Aldrich Chemie GmbH, Buchs, Switzerland) was assessed as previously described.^33^ Briefly, BLECs were washed twice and 50 μM of LY in permeability assay medium (PAM) (95% 1X HBSS +Ca/+Mg, 2.5% HEPES, 2.5% FBS) was added to the top wells. Filter inserts were then transferred every 20 min to wells containing fresh PAM over a total of 60 min at 37°C (5% CO2). Fluorescence was quantified at 432/538 nm (excitation and emission) setting using multiplate reader (Tecan Infinite M1000) and the permeability coefficient was calculated using the clearance principle as described previously.^32^ Three empty inserts were used as control.

Trans-endothelial electrical resistance (TEER) of monolayers formed by BLECs was assessed by impedance TEER measurements using the cellZscope2 device (Nanoanalytics, Muenster, Germany) as described previously.^34^ TEER was monitored over a total period of 48 hours at 37°C (8% CO2).

### T-cell binding assays under static conditions

T-cell binding assays to recombinant BBB cell-adhesion molecules under static conditions were performed as previously described.^35^ In brief, Teflon (PTFE) slides (Thermofisher Scientific) were coated with 10 μg/mL of either rhVCAM-1 (Biolegend), rhJAM-B (R&D system), hrfibronectin (R&D system) or hrMadCAM-1 (R&D system) in DPBS for 1 hour at 37°C and blocked with 1.5% (v/v) bovine serum albumin (Sigma-Aldrich) in DPBS over night at 4°C. Recombinant Delta/Notch-like EGF-related receptor (R&D system) which does not bind Notch was employed as negative control to test for unspecific T-cell interactions.

CMFDA pre-labelled human CD4 T cells, CD4 Th cells (Th1, Th2, Th1*, and Th17) or Treg cells were let to adhere to the immobilized recombinant adhesion molecules for 30 min at RT with gentle shacking. Slides were then washed twice with PBS and fixed with 2.5% (v/v) glutaraldehyde in PBS for 2 hours on ice. Adhered T cells were imaged using a Nikon Eclipse E600 microscope connected to a Nikon Digital Camera DXM1200F with Nikon NIS-Elements BR3.10 software (Nikon, Egg, Switzerland) and counted using Image J software (NIH, Bethesda, MD, USA).

### T-cell binding assays under physiological flow conditions

For live-cell imaging of human CD4 T-cell interaction with recombinant BBB cell-adhesion molecules under physiological flow conditions, µ-Dishes (35 mm, low, iBidi) were coated with 1,54 µg/mL rhVCAM-1 (Biolegend) and/or 1,14 µg/mL rhICAM-1 (R&D system) as described above. CMFDA-pre-labelled T cells were perfused on top of the pre-coated dishes at a concentration of 1 × 10^6^ cells/mL in migration assay medium (MAM) (DMEM w/o phenol red (Gibco), 5% (v/v) heat inactivated FBS (Hyclone), 4 mM L-Glutamine (Gibco), 25 mM HEPES (Gibco)) as previously described.^36^ In brief, a parallel flow chamber connected to an automated syringe pump (Harvard Apparatus, Holliston, MA, USA) was mounted on the pre-coated dishes and placed on the heating stage of an inverted microscope. T cells were then allowed to accumulate for 4 min at low shear stress (0.1 dyn/cm^2^) and their interaction with VCAM-1 and/or ICAM-1 was imaged under physiological shear stress (1.5 dyn/cm^2^) using an AxioObserver Z1 microscope (Carl Zeiss, Feldbach, Switzerland) connected to a digital camera (Carl Zeiss). Time-lapse videos were created by taking one image every 10 s over a 10 min period using the ZEN blue software (AxioVision, Carl Zeiss). As control, some CD4 T cells were incubated with 1 μg/mL of natalizumab (NTZ, Tysabri, Biogen) or 1 μg/mL of an anti-β2-integrins antibody (TS1/18, Invitrogen) for 30 min at 37°C (8% CO2) prior imaging. Off-line video analysis allowing to define the number of arrested T cells and their post-arrest behavior (e.g., mean crawling speed, distance, Euclidian distance, and directionality) was performed using Image J (NIH, Bethesda, MD, USA) and the Chemotaxis and Migration Tool software (iBidi). Directionality of T-cell crawling on VCAM-1 and ICAM-1 under physiological shear stress was calculated as forward migration index towards the x-axis (xFMI), where xFMI is the straight x-axis distance (Dx) covered by the T cells divided by the accumulated total distance (Dacc) of Th-cell movement (xFMI = Dx / Dacc).

### T-cell transmigration assays under static conditions

The ability of activated CD4 T cells to migrate across BLECs, as a human *in vitro* model of the BBB, was investigated as previously reported.^31,32^ In brief, BLECs were grown to confluency on 3.0μm pore Transwell^@^ filters. 2x10^5^ Cell Tracker^TM^ Green CMFDA labeled CD4 T cells were added to in the upper chamber on the BLEC monolayer and allowed to migrate across the BLEC monolayer for 8 hours at 37°C (10% CO2). CD4 T cells that had migrated across the BLEC monolayer as well as input cells were assessed by flow cytometry (Attune NxT Flow Cytometer, Thermofisher Scientific, Switzerland). Antibodies are listed in **Supplementary Table 8.**

### T-cell transmigration assays under physiological flow

BLECs were grown to confluency in μSiM-CVB flow chambers as described above. CMFDA prelabelled human CD4 T cells, Th1 or Treg were perfused on top of BLECs at a concentration of 1 × 10^6^ cells/mL in MAM as previously described.^33,35,37^ In brief, flow was applied by connecting each μSiM-CVB flow chamber to an automatic syringe pump (Harvard Apparatus, Holliston, MA, USA) on the heating stage of an inverted microscope. T cells were then allowed to accumulate for 4 min at low shear stress (0.1 dyn/cm^2^) and their interaction with BLECs was imaged under physiological shear stress (1.5 dyn/cm^2^) using an AxioObserver Z1 microscope (Carl Zeiss, Feldbach, Switzerland) connected to a digital camera (AxioVision, Carl Zeiss). As additional controls, some T cells were incubated with 1 μg/mL anti-α4-integrin antibody natalizumab (NTZ, Tysabri, Biogen), and 1 μg/mL of an anti-β2-integrins antibody (TS1/18, Invitrogen) for 30 min at 37°C (8% CO2) prior imaging. Time-lapse videos were created by taking one image every 10 s over a 24 min period using the ZEN blue software (Carl Zeiss). Off-line video analysis allowing to define the arrest of T cells on BLECs and their post-arrest behavior e.g., probing, crawling, and diapedesis, was performed using Image J software (NIH, Bethesda, MD, USA) as described before.^35,37^ The whole field of view (FOV) of the μSim-CVB devices (2 mm x 0.7 mm) was considered for the analysis. T cells crawling out of the FOV during image acquisition were not included in the behavioral analysis. Off-line video analysis of the mean crawling speed, the mean crawling distance, and the mean crawling Euclidian distance of CD4 T cells on BLECs was performed using Image J and Chemotaxis and Migration Tool (iBidi). Only T cells that were classified as crawling cells and that were followed over the full recording time were taken into consideration.

### Experimental autoimmune encephalomyelitis

EAE was induced in 8-12 weeks old female VE-Cadherin-GFP knock-in C57BL/6J mice as previously described.^38,39^ Pertussis toxin (List Biological Laboratories, Campbell, US) (300 ng in 100 μL PBS/mouse) was injected intraperitoneally (i.p.) on day 0 (immunization day) and day 2 post-immunization. Weights and clinical severity were assessed twice daily and scored as in^40^: 0, asymptomatic; 0.5, limb tail; 1, hind leg weakness; 2, hind leg paraplegia; 3, hind leg paraplegia and incontinence.

### 2-photon intravital microscopy of the cervical spinal cord

2-photon intravital microscopy of cervical spinal cord of VE-Cadherin-GFP knock-in C57BL/6J mice was performed as previously described.^41^ In brief, mice were anesthetized and fixed to a stereotaxic frame for cervical spinal cord surgery. Mice were transferred to the TrimScope-II 2-photon microscope system equipped with an Olympus BX51WI fluorescence microscope (LaVision BioTec) and imaged with a laser wavelength of 890 nm. Afterwards, four FOVs, two on each side of the central posterior dorsal vein, with width (x) and height (y) of 443 µm and a depth (z) of 100 µm were chosen. The FOV was selected for vessels approximately at the second to third generation of branching away from the central posterior dorsal vein, considered as post capillary venules. 3D images were acquired every 2 min over a total period of 1 hour. Pre-labelled (CMFDA or cell tracker Deep Red) 1,25(OH)2D3 and vehicle control treated human CD4 T cells were co-injected (5 × 10^6^ each) systemically via a carotid artery catheter 4 min after starting imaging.^41^ Distortion correction of the images during 2-photon-imaging was performed by using Vivo Follow 2.0.^42^ Imaris 9.8 software (Oxford Instruments Group) was used to transform sequences of image stacks into volume-rendered 4D images and analyze the arrest and post-arrest interactions of CD4 T cells with the endothelium.

### RNAseq data analysis

RNAseq data reanalysed for this study were taken from the dataset deposited at the Gene Expression Omnibus (GEO) under GSE154741.^16^

### Statistical analysis

Data are shown as the mean ± SD or SEM. Statistical significance between two groups was assessed by unpaired or paired t-test, while the comparison between within one or multiple groups and conditions was assessed by either One-or Two-way ANOVA followed by Šídák’s multiple comparisons test. Precise statistical analysis and relative significance are indicated in the corresponding figures and figure legends (p < 0.05 = *, p < 0.01 = **, p < 0.001 = ***, p < 0.0001 = ****). Statistical analyses comprising calculation of degrees of freedom were performed using GraphPad Prism 9 software (Graphpad software, La Jolla, CA, USA).

## Results

### Vitamin D reduces the cell surface expression of cell adhesion molecules on CD4 T cells

To understand the potential effect of VitD on T-cell migration across the BBB, we first determined the effect of the active form of VitD, namely 1,25-Dihydroxyvitamin D3 (1,25(OH)2D3), from now on referred to as VitD, on the expression levels of a wide panel of adhesion molecules involved in T-cell migration across the BBB. Human CD4 T cells were *in vitro* activated in the presence of 100nM of VitD or vehicle control (EtOH) prior to analysis by flow cytometry. Activated CD4 T cells showed homogenous cell surface expression of α4, αL-, β2-, α5- and α6-integrin subunits, while immunostaining for β1- and β7-integrin subunits and PSGL-1 distinguished two cell subsets of CD4 T cells with high and low expression levels, respectively **(Figure 1).** VitD significantly reduced cell surface expression (MFI) of all adhesion molecules on CD4 T cells compared to vehicle control with the exception to the β1-integrin^low^, β7-integrin^low^ and PSGL-1^high^ subsets which remained unaltered and α5-integrins which were increased **(Figure 1).** While VitD did not change the percentage of β1-integrin^high^ and β1-integrin^low^ CD4 T cells **(Figure 1),** it significantly reduced the β7-integrin^high^ and PSGL-1^high^ CD4 T cell subsets compared to a significant increase in the β7-integrin^low^ and PSGL-1^low^ CD4 T cell subsets **(Figure 1 F, H).** Re-analyzing previously published RNA-seq data of CD3/CD28-activated human CD4 T cells in the presence or absence of VitD (GEO GSE154741) ^16^ underscored that VitD reduced expression levels of α4, αL-, β2-integrins also at the mRNA level with the exception to β2-integrins (**Supplementary Fig. 1A**).

**Figure 1.**
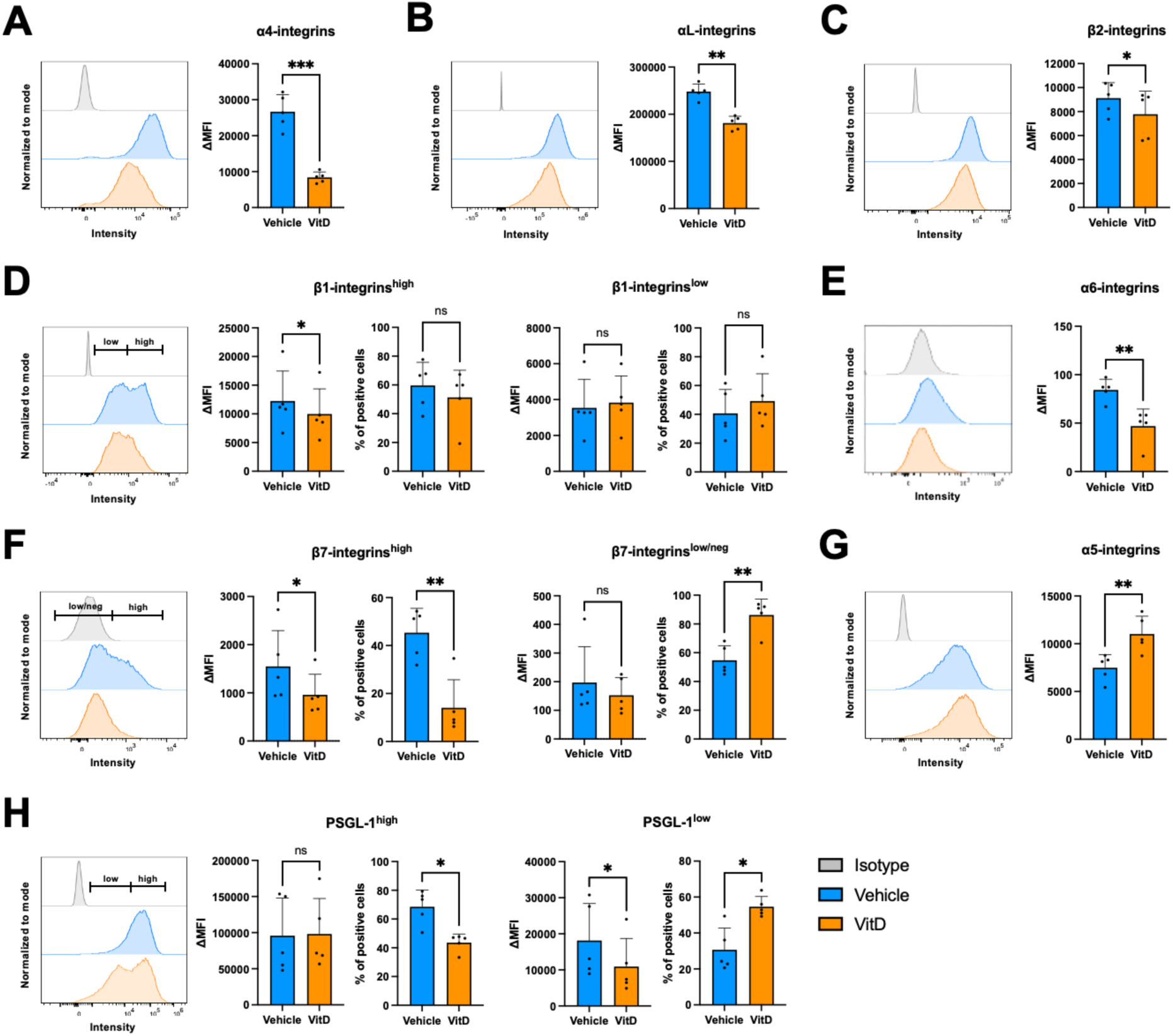
Vitamin D reduces the cell-surface expression of important integrins for the migration of CD4^+^ T cells across the BBB. (A-H) Representative histogram plots, geometric ΔMFI (MFI specific staining - MFI isotype) and percentage of positive cells for the cell-surface expression of different adhesion molecules on CD4^+^ T cells treated with 100nM of 1,25(OH)2D3 (orange) or vehicle control (blue). Isotype control for each flow cytometry staining is shown in grey in each histogram plot. Bar graphs show the mean ± SD of 5 independent experiments for the cell-surface expression analysis of α4-integrins **(A)**, αL-integrins **(B)**, β2-integrins **(C)**, β1-integrins **(D)**, α6-integrins **(E)**, β7-integrins **(F)**, α5-integrins **(G)**, and PSGL-1 **(H)**. **(D, F, H)** When two distinct population were identified, CD4^+^ T cells were subdivided in cells expressing high and low or low/neg cell-surface levels of the different adhesion molecules for analysis. The gating strategy for high and low or low/neg populations is displayed in each corresponding histogram plot. Statistical analysis: paired t-test (p < 0.05 = *, p < 0.01 = **, p < 0.001 = ***, p < 0.0001 = ****).

We also found that VitD modulated additional adhesion molecules by increasing CD4 T cell surface levels of platelet endothelial cell-adhesion molecule 1 (PECAM-1) and decreasing those of L-selectin while leaving CD99 cell surface levels unaltered **(Supplementary Fig. 1)**. In addition, VitD increased and reduced the percentage of PECAM-1^high^ and L-selectin^high^ CD4 T cells, respectively and significantly reduced activated leukocyte cell adhesion molecule (ALCAM) **(Supplementary Fig. 1)**.

Taken together, our results show that VitD modulates the cell surface expression of several key adhesion molecules on CD4 T cells. Importantly, VitD reduced the T-cell surface levels of α4-and β2-integrins, crucial integrins molecules involved in T-cell migration across the BBB.^22^ These observations suggest that VitD may reduce CD4 T-cell migration across the BBB.

### Vitamin D decreases cell surface adhesion molecule expression on effector but not on regulatory CD4 T cells

We next asked if VitD differentially affects adhesion molecule expression on distinct Th subsets (Th1, Th1*, Th2, Th17 and FoxP3^+^ CD4 Treg) **(Supplementary Figs. 2 and 3**). All CD4 T-cell subsets isolated from the blood of healthy donors showed a homogenous – single peak - cell surface expression of α4-, α5-, α6-, αL-, β2-integrin subunits and PSGL-1, while two distinct sub-populations with high and low or low/negative were observed for β1- and β7-integrins **(Figure 2).** VitD significantly reduced α4- and β2-integrins on Th1, Th1*, Th17 and Th2 cells but not on Treg compared to vehicle control. All CD4 T-cell subsets showed a VitD-reduced cell surface expression of αL-and β7-integrins. However, VitD significantly reduced β1-integrins only on Th17 cells and α5-integrins only on Treg and the levels of PSGL-1 only on Th1 and Th1* cells. Despite very low surface expression of α6-integrins on all CD4 T-cell subsets, VitD significantly decreased α6-integrins levels of Th17 cells. These data show that VitD specifically decreases the cell surface expression of those adhesion molecules, especially α4- and β2-integrins involved in T-cell migration across the BBB rather on Teff than Treg. VitD might thus selectively reduce the migration of Teff but not Treg across the BBB.

**Figure 2.**
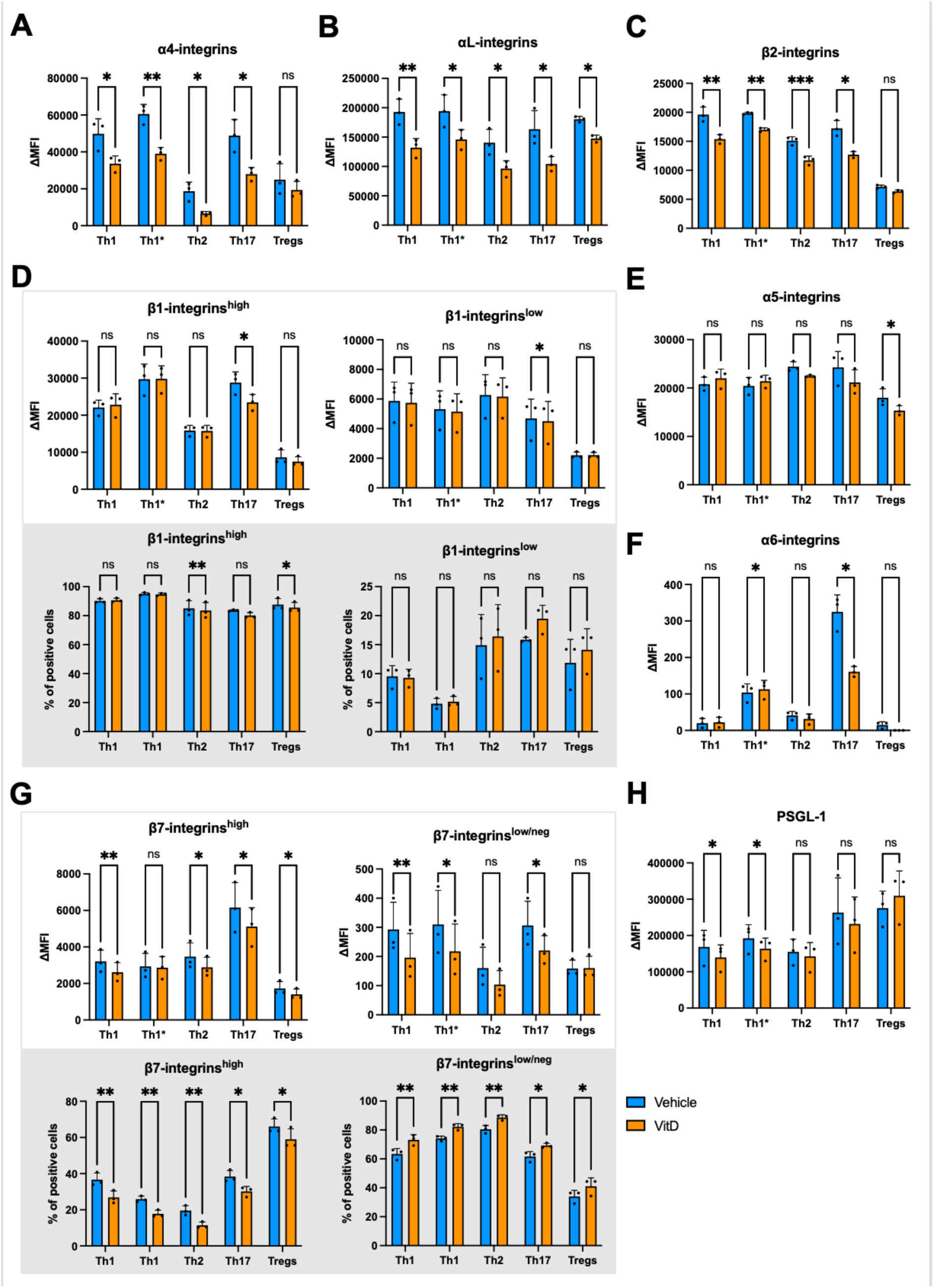
Vitamin D differentially modulates surface expression of key integrins for the migration across the BBB in the different CD4^+^ T-cell subsets. (A-H) Geometric ΔMFI (MFI specific staining - MFI isotype) and percentage of positive cells for the cell-surface expression of different adhesion molecules on different CD4^+^ T-cell subsets (Th1, Th1*, Th2, Th17, and Treg) of 3 healthy donors treated with 100nM of 1,25(OH)2D3 (orange) or vehicle control (blue). Bar graphs show the mean ± SD of 3 independent experiments for the cell-surface expression analysis of α4-integrins **(A)**, αL-integrins **(B)**, β2-integrins **(C)**, β1-integrins **(D)**, α5-integrins **(E)**, α6-integrins **(F)**, β7-integrins **(G**), and PSGL-1 **(H)**. **(D, G)** When two distinct population were identified, CD4^+^ T-cell subsets were subdivided in cells expressing high and low or low/neg cell-surface levels of the different adhesion molecules for analysis. Statistical analysis: paired t-test (p < 0.05 = *, p < 0.01 = **, p < 0.001 = ***, p < 0.0001 = ****).

**Figure 3.**
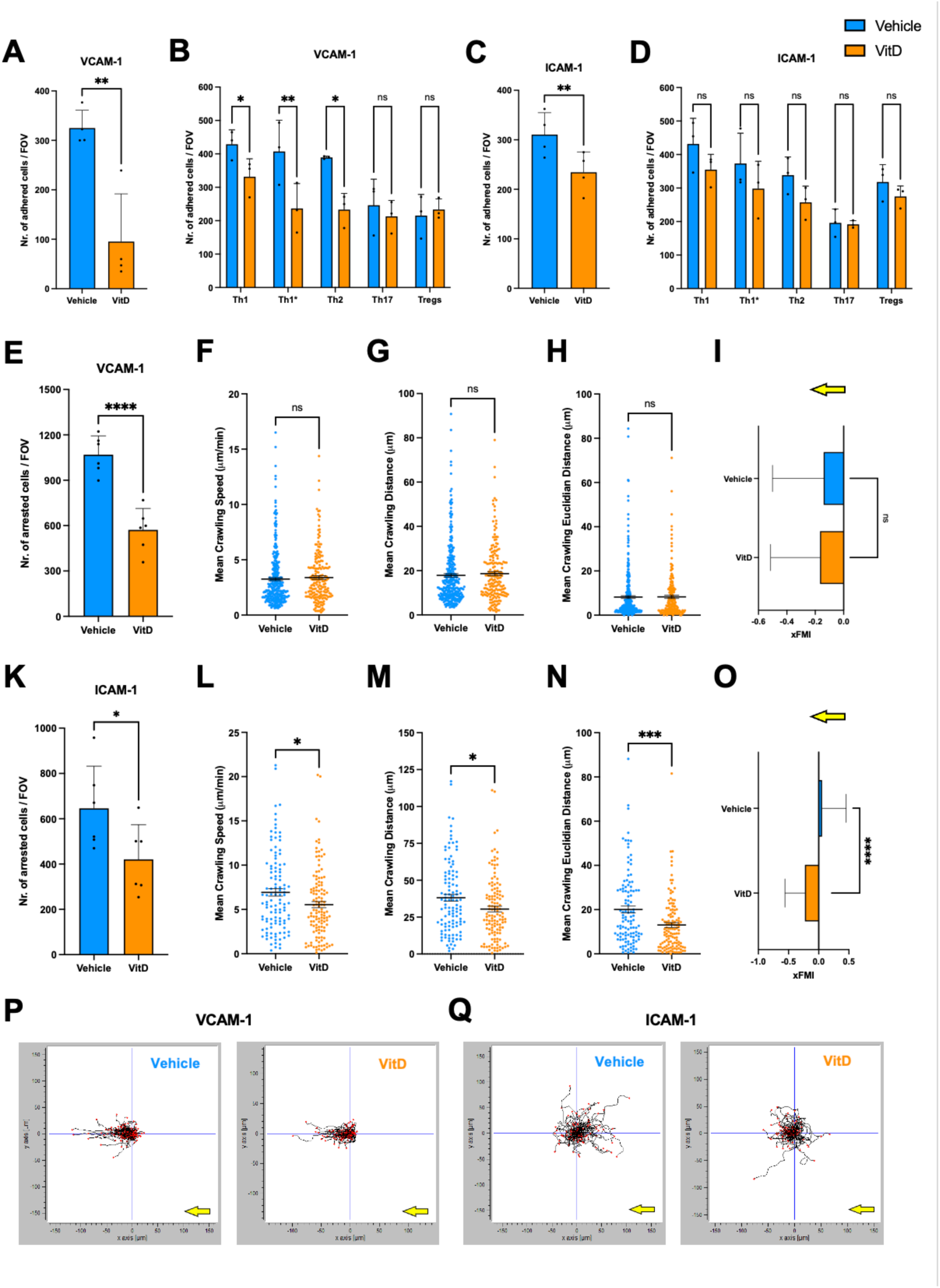
Vitamin D impacts adhesion and post-arrest behavior of T cells on VCAM-1 and ICAM-1. Number of adhered human CD4^+^ T cells **(A, C)** and different CD4^+^ Th subsets (Th1, Th1*, Th2, Th17, and Treg) **(B, D)** treated with 100nM of 1,25(OH)2D3 (orange) or vehicle control (blue) to immobilized recombinant VCAM-1 **(A, B)**, ICAM-1 **(C,D)** under static conditions. Bar graphs show the mean ± SD of 3-4 independent experiments done in triplicates. Statistical analysis: paired t-test (p < 0.05 = *, p < 0.01 = **, p < 0.001 = ***, p < 0.0001 = ****). **(E,K)** Number of arrested human CD4^+^ T cells treated with 100nM of 1,25(OH)2D3 (orange) or vehicle control (blue) to equimolar concentrations of immobilized recombinant VCAM-1 (1.54 μg/mL) **(E)** and ICAM-1 (1.14 μg/mL) **(K)** under physiological flow conditions. Bar graphs show the mean ± SD of 6 movies of 2 independent experiments. Mean crawling speed **(F, L)**, distance **(G, M)** and Euclidian distance **(H, N)** of CD4^+^ T cells on VCAM-1 **(E-H)** and ICAM-1 **(K-N)** under physiological flow conditions. Graphs show the mean ± SEM for 1 representative experiment in each group (>100 cells per group were analysed). Each data point represents the velocity, the distance, and the Euclidian distance of one cell. **(I, O)** Mean directionality of CD4^+^ T-cell crawling on VCAM-1 **(I)** and ICAM-1 **(O)** under physiological flow conditions. Bar graphs show the mean ± SEM of T-cell crawling directionality expressed as xFMI (xFMI = Dx/Dacc, Dx: straight x-axis distance covered by the T cell, Dacc: accumulated total distance of T-cell movement). **(P, Q)** x/y diagrams of CD4^+^ T-cell crawling tracks on VCAM-1 **(P)** and ICAM-1 **(Q)** under physiological flow. Vehicle control and 1,25(OH)2D3 treatment conditions are shown on the left and on the right respectively. For each track, the site of arrest was set to the center point of the respective diagram (shown by the intersection of two blue lines). End points of tracks are indicated by a red dot. Flow direction is illustrated by a yellow arrow and was along the x-axis from plus to minus **(I, O-Q).** Statistical analysis: **(E, K)** unpaired t-test and **(F-I, L-O)** Mann-Whitney test (p < 0.05 = *, p < 0.01 = **, p < 0.001 = ***, p < 0.0001 = ****).

### Vitamin D reduces effector but not regulatory CD4 T cell adhesion to VCAM-1

To explore the functional impact of VitD-reduced cell surface expression of α4- and β2-integrins, we next investigated the adhesion of VitD-treated CD4 T cells to immobilized rVCAM-1 and rICAM-1 under static conditions. Correlating with VitD-reduced cell surface expression of α4β1- and αLβ2- integrins, VitD treatment significantly reduced the adhesion of CD4 cells to rVCAM-1 and rICAM-1 **(Figure 3A, 3C).**

When exploring the impact of VitD on binding of the different Th subsets to VCAM-1 and ICAM-1, we observed that although VitD reduced cell surface expression of α4β1-integrins on all CD4 Teff-subsets **(Figure 2),** it only significantly reduced adhesion of Th1, Th1* and Th2 cells binding in high numbers to VCAM-1 but not of Th17 cells which showed lower binding to VCAM-1 **(Figure 3B)**. In accordance with the lack of an effect of VitD on α4β1-integrin expression on Treg, it also did not affect binding of Treg to VCAM-1 (**Figure 3B**). Although VitD treatment induced a trend towards lower binding of Th1, Th1* and Th2 cells to ICAM-1, we observed no significant effect on ICAM-1 binding of Th17 and Treg (**Figure 3D**). These data show that VitD significantly reduces the binding of potentially pathogenic Th1 and Th1* but not of Treg to VCAM-1, suggesting that VitD may favor the arrest of Treg over Teff on the inflamed BBB endothelium.

### Vitamin D impacts CD4 T cell interaction with ICAM-1 and VCAM-1 under physiological flow *in vitro*

It has previously been shown that, under physiological flow, endothelial VCAM-1 mediates arrest of T cells on the BBB and endothelial ICAM-1 mediates post-arrest T cell crawling against the direction of blood flow to sites of diapedesis.^43^ We therefore examined the effect of VitD on CD4 T-cell interaction dynamics with equimolar concentrations of immobilized rVCAM-1 or rICAM-1 under physiological flow by in vitro live-cell imaging. CD4 T cells readily arrested to VCAM-1 and ICAM-1 **(Figure 3)**. In accordance with our observations under static conditions, VitD significantly reduced shear-resistant arrest of CD4 T cells to rVCAM-1 and IrCAM-1 compared to vehicle control (**Figure 3E,K)**. Shear-resistant arrest of CD4 T cells to VCAM-1 and ICAM-1 was mediated by α4- and β2-integrins, respectively, as confirmed by abolishing CD4 T cells arrest with the respective integrin-blocking antibodies **(Supplementary Movie 1, 2).**

By analysing the post-arrest behaviour of CD4 T cells on VCAM-1 and ICAM-1, we observed that VitD significantly influenced post-arrest behavior of CD4 T cells on ICAM-1 but not VCAM-1 **(Figure 3E-O).** VitD reduced the mean crawling speed, distance, and Euclidian distance of CD4 T cells on ICAM-1 under physiological flow **(Figure 3L-O)**. In accordance with previous observations^35,36^, we found that CD4 T cells crawled in the direction of flow on VCAM-1 and preferentially crawled against the direction of flow on ICAM-1 **(Figure 3 I,P,O, Q; Supplementary Movie 3)**. Interestingly, VitD significantly reduced the ability of CD4 T cells to crawl against the direction of the flow on ICAM-1 **(Figure 3)**. Taken together these data show that VitD reduces shear resistant arrest of CD4 T cells to VCAM-1 and ICAM-1 and impairs post-arrest T-cell crawling against the direction of flow on ICAM-1 thus leading to reduced T cell diapedesis across the BBB.

### Vitamin D modulates CD4 T cell arrest and post-arrest behaviour on the BBB under physiological flow *in vitro*

As shear stress critically influences the migration of T cells across the BBB^36,44^, we next assessed the effect of VitD on the arrest and the post-arrest behavior of CD4 T cells on pro-inflammatory cytokine (TNFα + IFNγ) stimulated BLECs as an *in vitro* model of the human BBB, under physiological flow by *in vitro* live-cell imaging. VitD significantly reduced the arrest of CD4 T cells on BLECs compared to vehicle control **(Figure 4A)**. Antibody-mediated blocking of α4- and β2-integrins on CD4 T cells completely abrogated CD4 T-cell arrest on BLECs **(Supplementary Movie 4)**, underscoring the essential roles of α4β1- and αLβ2-integrins in mediating shear-resistant arrest of T cells on the BBB. Furthermore, VitD significantly reduced CD4 T cell probing followed by diapedesis leading to increased numbers of T cells that remained crawling on the BLEC monolayer **(Figure 4B)**. Interestingly, although only a minor fraction of CD4 T cells detached from the BLEC monolayer during the observation time, this was further reduced after VitD treatment. Despite the significant effects of VitD on CD4 T cell interaction with BLECs under flow, VitD did not significantly reduce the overall CD4 T-cell diapedesis across BLECs under physiological flow **(Figure 4C)**. However, VitD decreased the mean crawling speed, distance, and Euclidian distance of CD4 T cells crawling on BLECs **(Figure 4D-F; H, I)** without affecting their crawling directionality (xFMI) on the BLEC monolayer **(Figure 4G; Supplementary Movie 5).** This further underscores a role for VitD in decreasing integrin activation leading to insufficient traction.

**Figure 4.**
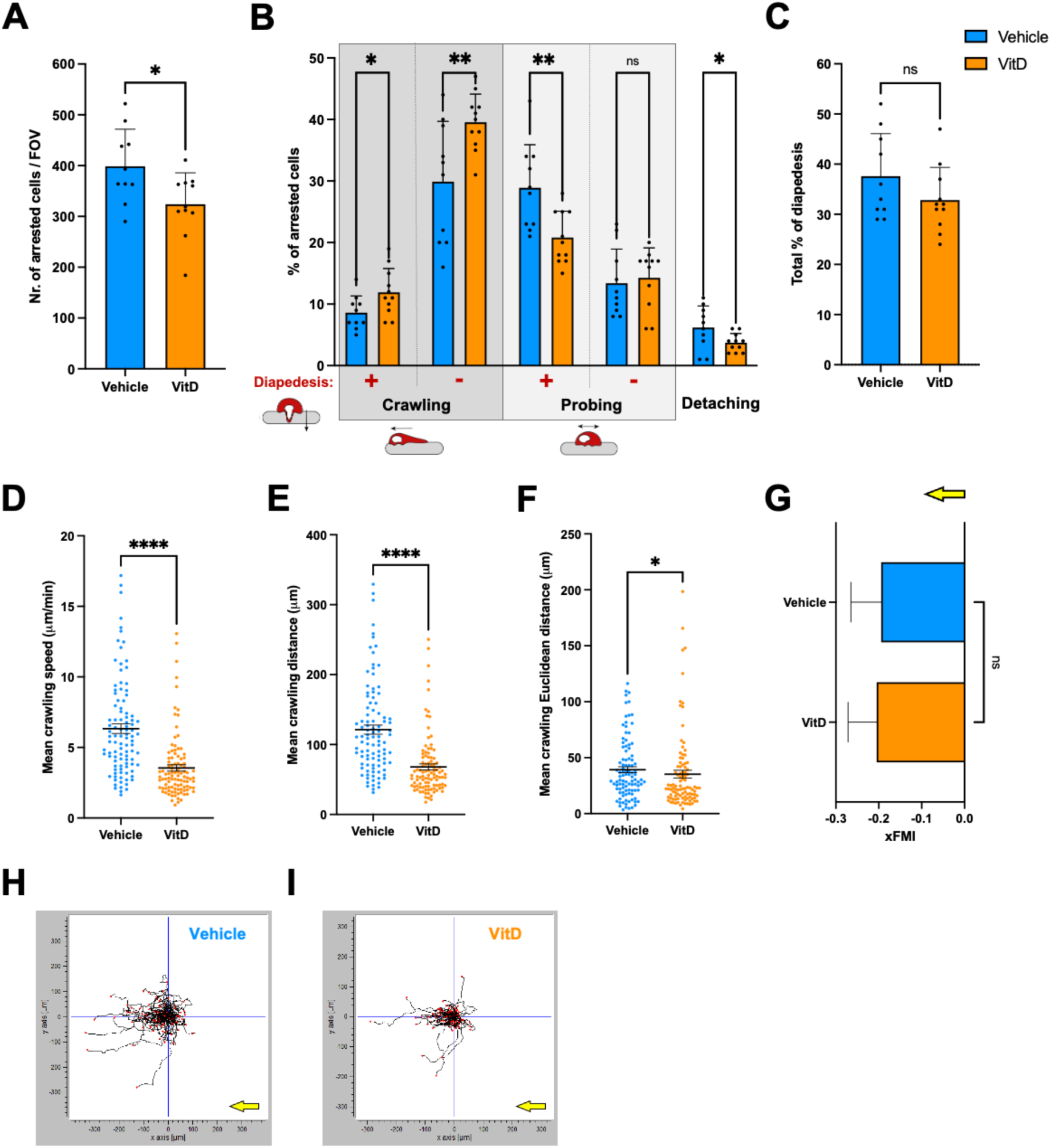
Vitamin D modulates CD4^+^ T-cell arrest and post-arrest behavior on the BBB under flow *in vitro*. **(A)** Number of arrested human CD4^+^ T cells treated with 100nM of 1,25(OH)2D3 (orange) or vehicle control (blue) to 16-24 h pro-inflammatory cytokine (76 IU/mL TNFα + 20 IU/mL IFNγ) stimulated brain-like endothelial cells (BLECs) under flow conditions. **(B)** Quantification of the post-arrest dynamic behavior of CD4^+^ T cells on stimulated BLECs under physiological flow conditions during 20 min of recording. Behavioral categories were subdivided in crawling, probing, and detached cells and are shown as fraction of all shear resistant arrested CD4^+^ T cells (100%). Within the categories probing and crawling we additionally distinguished the categories with and without of subsequent diapedesis. **(C)** Overall percentage of CD4^+^ T-cell diapedesis across BLECs. **(A-C)** Bar graphs show mean ± SD for 10-11 movies of 3 independent experiments. Mean crawling speed **(D)**, distance **(E)** and Euclidian distance **(F)** of CD4^+^ T cells on stimulated BLECs under physiological flow conditions. Graphs show mean ± SEM of >100 cells per treatment group. Each data point represents the velocity, the distance, and the Euclidian distance of one cell. **(G)** Directionality of CD4^+^ T-cell crawling on stimulated BLECs under physiological flow conditions. Graph bars show mean ± SEM expressed as xFMI (xFMI = Dx/Dacc, Dx: straight x-axis distance covered by the T cell, Dacc: accumulated total distance of T-cell movement). The direction of the physiological flow is indicated by a yellow arrow and was along the x-axis from plus to minus. Statistical analysis: **(A-C)** unpaired t-test and **(D-G)** Mann-Whitney test (p < 0.05 = *, p < 0.01 = **, p < 0.001 = ***, p < 0.0001 = ****). **(H, I)** x/y diagrams of human CD4^+^ T-cell crawling tracks on 16-24 h pro-inflammatory cytokine (76 IU/mL TNFα + 20 IU/mL IFNγ) stimulated BLECs under physiological flow. Vehicle control **(H)** and 1,25(OH)2D3 treatment **(I)** conditions are shown. For each track, the site of arrest was set to the center point of the respective diagram (shown by the intersection of two blue lines). End points of tracks are indicated by a red dot. Flow direction is illustrated by an arrow (yellow).

### Vitamin D differentially impacts the migration of effector and regulatory T cells across the BBB under static and physiological flow conditions *in vitro*

Based on the differential effects of VitD on adhesion molecule expression, we next assessed whether VitD differentially affects the diapedesis of Treg versus Teff across pro-inflammatory cytokine (TNFα + IFNγ)-stimulated BLECs first under static conditions. CD4 T cells preactivated in the presence or absence of VitD were assessed and the percentage of CD25^high^CD127^low^ Treg and CD127^high^ Teff before and after diapedesis across BLEC monolayers was determined. VitD increased the percentage of transmigrated Treg, while it decreased the percentage of Teff, with a three-fold change in the Treg/Teff ratio before and after diapedesis across BLECs **(Figure 5A)**. These data further underscore that VitD favors the migration of Treg over Teff across the BBB *in vitro*.

**Figure 5.**
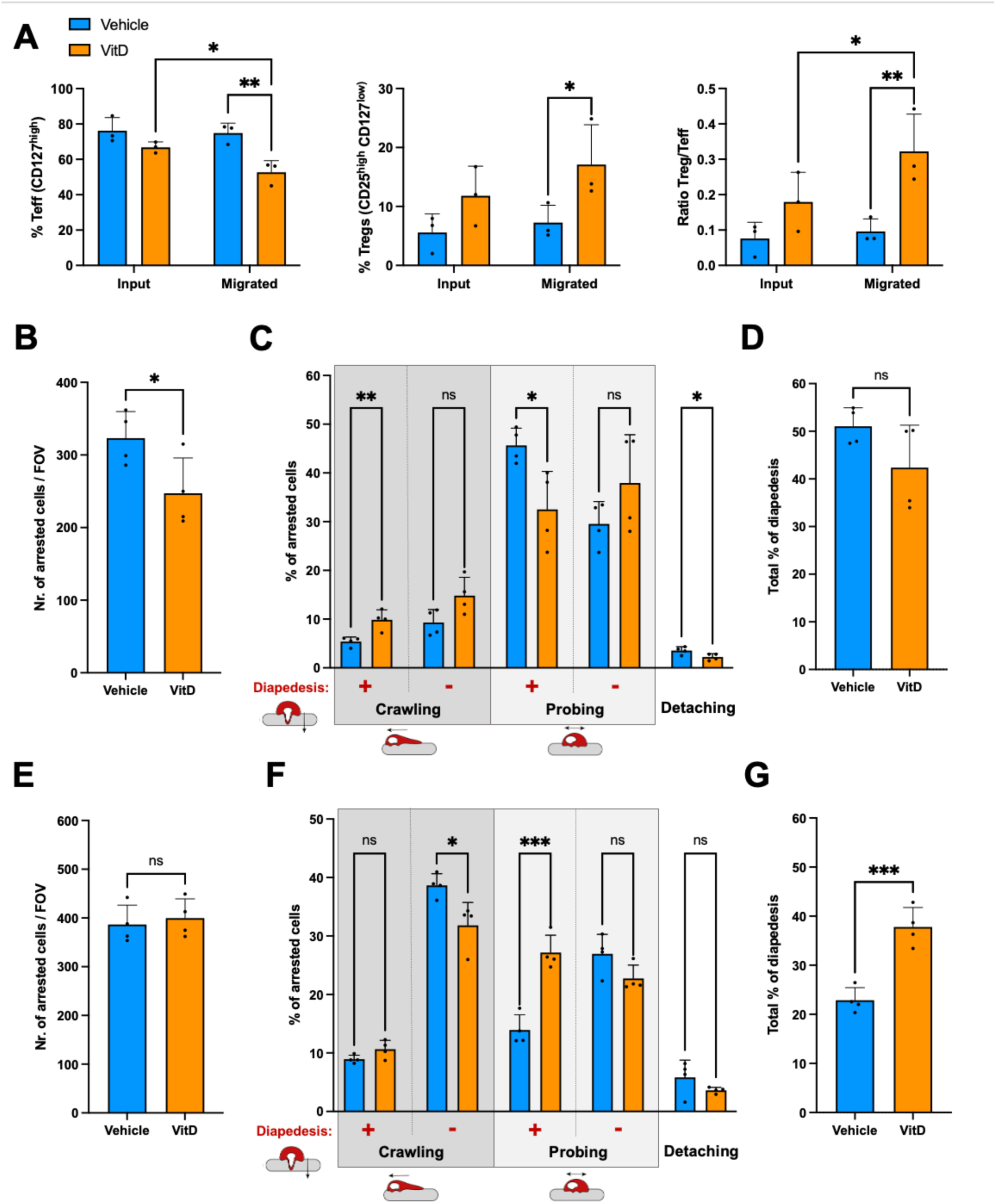
Vitamin D favours the migration of regulatory over effector CD4^+^ T cells across the BBB *in vitro*. **(A)** Analysis of the percentages of regulatory (CD25^high^CD127^low^) and effector (CD127^high^) CD4^+^ T cells prior to (input) and after migration across brain-like endothelial cells (BLECs) monolayers (migrated). The Treg/Teff ratio is shown on the right. Bar graphs show mean ± SD of 3 independent experiments done in duplicates or triplicates. Number of arrested human Th1 **(B)** and Treg **(E)** treated with 100nM of 1,25(OH)2D3 (orange) or vehicle control (blue) to 16-24 h pro-inflammatory cytokine (76 IU/mL TNFα + 20 IU/mL IFNγ) stimulated BLECs under flow conditions. Quantification of the post-arrest dynamic behavior of Th1 **(C)** and Treg **(F)** on stimulated BLECs under physiological flow conditions during 20 min of recording. Behavioral categories were subdivided in crawling, probing, and detached cells and are shown as fraction of all shear resistant arrested T cells (100%). Within the categories probing and crawling we additionally distinguished the categories with and without of subsequent diapedesis. Overall percentage of Th1 **(D)** and Treg **(G)** cell diapedesis across BLECs. **(B-G)** Bar graphs show mean ± SD for 4 movies of 2 independent experiments. Statistical analysis: **(A)** One-way ANOVA followed by Šídák’s multiple comparisons test and **(B-G)** unpaired t-test (p < 0.05 = *, p < 0.01 = **, p < 0.001 = ***, p < 0.0001 = ****).

In order to gain a better understanding of the molecular mechanisms affected by VitD on the behaviour of the different T cell subsets on the inflamed BBB, we next investigated the interaction and migration of one key Teff subset, Th1, versus Treg across BLECs under physiological flow by *in vitro* live-cell imaging. While VitD significantly reduced the number of Th1 cells arresting on BLECs under physiological flow, it did not reduce arrest of Treg compared to vehicle control **(Figure 5B, E)**. Furthermore, VitD treatment differentially affected the post-arrest behavior, namely probing, crawling and diapedesis of Th1 and Treg on stimulated BLECs **(Figure 5C, F)**. More than 50% of arrested Th1 cells succeeded to cross the BLEC monolayer with the larger fraction after probing. In contrast the majority of Treg crawled on the BLEC surface with less than 20% successfully crossing the BLEC monolayer during the observation time. Treatment with VitD significantly reduced the fraction of probing Th1 cells undergoing diapedesis resulting in a significant reduction of Th1 diapedesis across the BLEC monolayer under flow **(Figure 5D)**. In sharp contrast, VitD significantly increased the fraction of Treg probing followed by diapedesis leading to a significant increase in the percentage of Treg undergoing diapedesis across the BLEC monolayer under physiological flow **(Figure 5G)**. Altogether, VitD reduced Th1- but not Treg arrest on BLECs under physiological flow and differentially affected post-arrest Th1- and Treg behaviour resulting in reduced diapedesis of Teff while promoting that of Treg across the BBB under physiological flow *in vitro*.

### Vitamin D does not stabilize BBB integrity or BBB immune quiescence *in vitro*

Endothelial cells including those of the BBB express the VDR suggesting that VitD deficiency can also lead to endothelial dysfunction.^45^ We therefore assessed the impact of VitD on the integrity and adhesion molecule profile of BLEC monolayers used in this study as *in vitro* model of the BBB.^32^ Immunofluorescence staining confirmed that VDR expression in BLECs in the nucleus and cytoplasm **(Figure 6A)**, allowing BLECs to respond to VitD treatment. Junctional localization of the tight- and adherens proteins claudin-5, ZO-1, and VE-cadherin remained unaltered upon VitD treatment of BLECs **(Figure 5A-D, L; Supplementary Fig. 4B-D)**.VitD did not modulate barrier integrity of BLEC monolayers as it failed to stabilize their cytokine-induced barrier dysfunction, as observed by a decline in TEER and an increase in permeability for the small molecular tracer lucifer yellow (LY) **(Figure 6E-G)**. Also, VitD did not modulate the cell surface expression of the adhesion molecules ICAM-1, VCAM-1, CD99, PECAM-1 and ICAM-2 on BLECs under both NS and stimulated conditions **(Figure 6 H-O, Supplementary Fig. 4)**. Altogether, our data show that VitD does not stabilize barrier functions nor immune quiescence of the BBB *in vitro* underscoring that VitD mediated alterations of T cell interaction with the BBB are due to altered behaviour of the T cell subsets.

**Figure 6.**
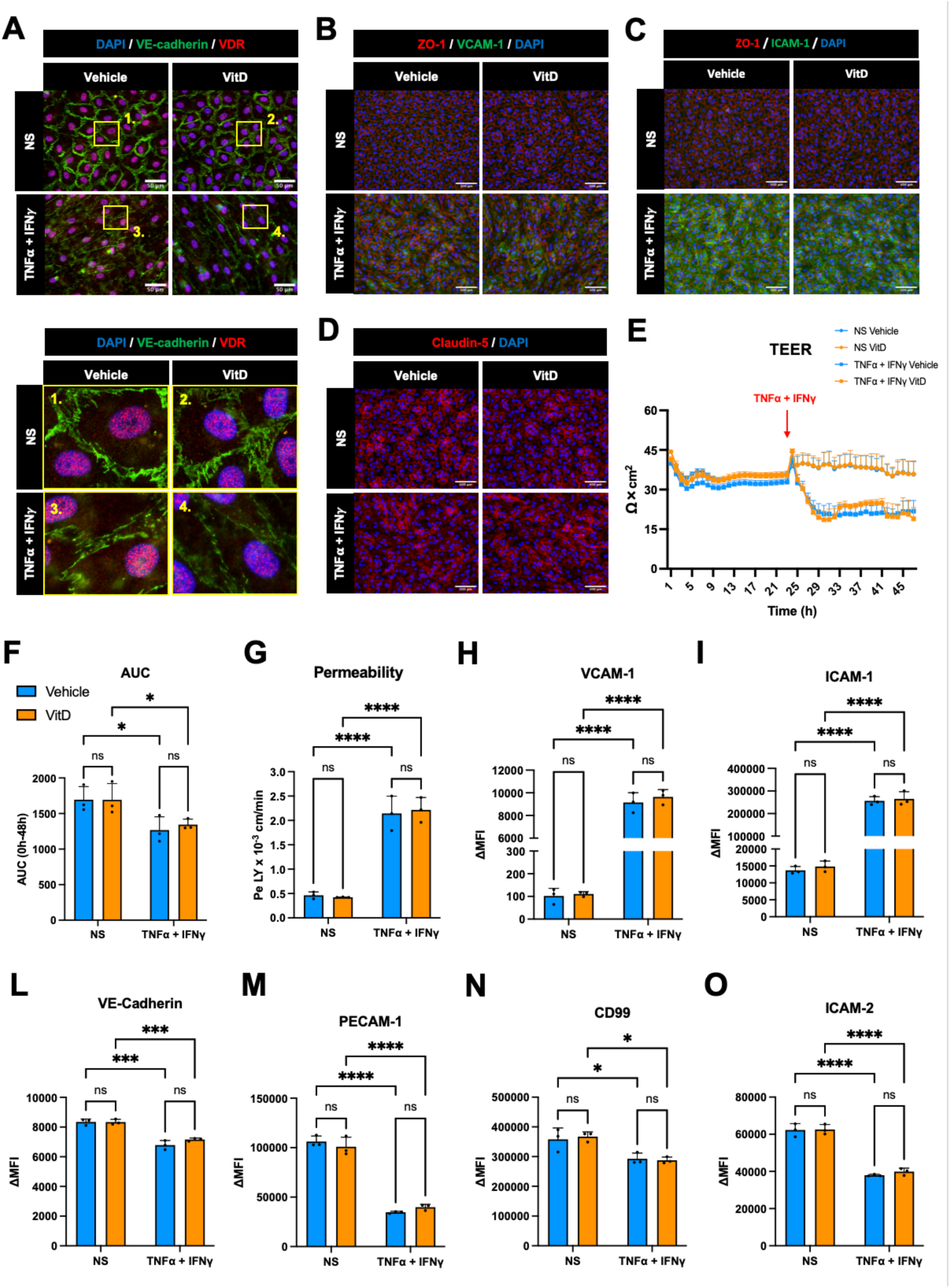
Vitamin D does not rescue BBB integrity nor upregulation of adhesion molecules upon inflammation *in vitro*. (A-D) Immunofluorescence staining of non-stimulated (NS) and 16 h pro-inflammatory cytokine stimulated (76 IU/mL TNFα + 20 IU/mL IFNγ) BLECs pre-treated for 2 d with 100nM of 1,25(OH)2D3 or vehicle control. **(A)** Immunostaining for VDR (red) and VE-cadherin (green) is shown. Scale bar = 50 μm. Zoom in or the yellow squares (1-4) is shown on the bottom. Immunostainings for VCAM-1 (green) **(B),** ICAM-1 (green) **(C),** and ZO-1 (red) are shown. Scale bar = 100 μm. Immunostaining for claudin-5 (red) **(D)** is shown. Scale bar = 100 μm. **(A-D)** Nuclei were stained with DAPI (blue). Data are representative of 3 independent experiments. **(E)** Time-dependent progression of the trans-endothelial electrical resistance (TEER) of BLEC monolayers over 48 hours. 1,25(OH)2D3 (orange) or vehicle control (blue) pre-treated BLECs were stimulated with pro-inflammatory cytokine (76 IU/mL TNFα + 20 IU/mL IFNγ, squares) or not (NS, circles) after 24 hours as indicated by the red arrow. Solid lines represent mean and error bars show ± SEM. **(F)** Quantification of TEER measurements. Bar graphs show the mean ± SD of the area under the curve (AUC) from 0 to 48 hours of TEER measurments. **(G)** BLECs permeability for 0.45 kDa Lucifer Yellow (LY). Bar graphs show the mean permeability coefficient (Pe ξ 10^-3^ cm/min) ± SD of diffused tracer across BLECs. Flow cytometry analysis for the cell-surface expression of VCAM-1 **(H)**, ICAM-1 **(I)**, VE-cadherin **(L)**, PECAM-1 **(M)**, CD99 **(N)** and ICAM-2 **(O)** on BLECs under non-stimulated (NS) and 16 h pro-inflammatory cytokine stimulated conditions (76 IU/mL TNFα + 20 IU/mL IFNγ) are shown. Bar graphs show geometric ΔMFI (MFI specific staining - MFI isotype) ± SD of 3 independent experiments. Statistical analysis: **(F-O)** Two-way ANOVA followed by Šídák’s multiple comparisons test (p < 0.05 = *, p < 0.01 = **, p < 0.001 = ***, p < 0.0001 = ****).

### Vitamin D reduces CD4 T-cell arrest and post-arrest behaviour on the inflamed BBB *in vivo*

We next assessed whether VitD affects the migration of T cells across the BBB *in vivo.* We studied the impact of VitD on the interaction of CD4 T cells with the inflamed BBB by intravital two-photon-microscopy *in vivo* in EAE. VitD-treated CD4 T cells and control-treated CD4 T cells were co-injected via a carotid artery catheter into VE-cadherin-GFP transgenic C57BL/6J mice suffering from EAE (d14-d17 post immunization, clinical score 0.5-1,5) and their interaction with spinal cord microvessels was observed using a cervical spinal cord window^41,42^. The CD4 T cell subsets were distinguished by cell tracker far red and CMFDA-labelling, respectively to avoid tracer effects **(Figure 7; Supplementary Fig. 5).** VitD pretreatment significantly reduced the arrest of CD4 T cells in inflamed cervical spinal cord microvessels compared to vehicle control **(Figure 7A,B)**. Analysing the post-arrest behaviour CD4 T cells in spinal cord microvessels did not show significant differences between VitD-and vehicle control-treated T cells with respect to traction of T cells crawling and probing **(Figure 7C)**. However, VitD-treated CD4 T cells showed a trend towards crawling in lower numbers against the direction of the blood flow **(Figure 7C)**. Furthermore, VitD significantly decreased the crawling speed of CD4 T cells in inflamed spinal cord microvessels **(Figure 7D)**. Thus, in accordance with our *in vitro* observations, VitD reduced CD4 T-cell arrest to and crawling speed on the inflamed BBB *in vivo*, suggesting that VitD reduces T cell entry across the BBB into the CNS.

**Figure 7.**
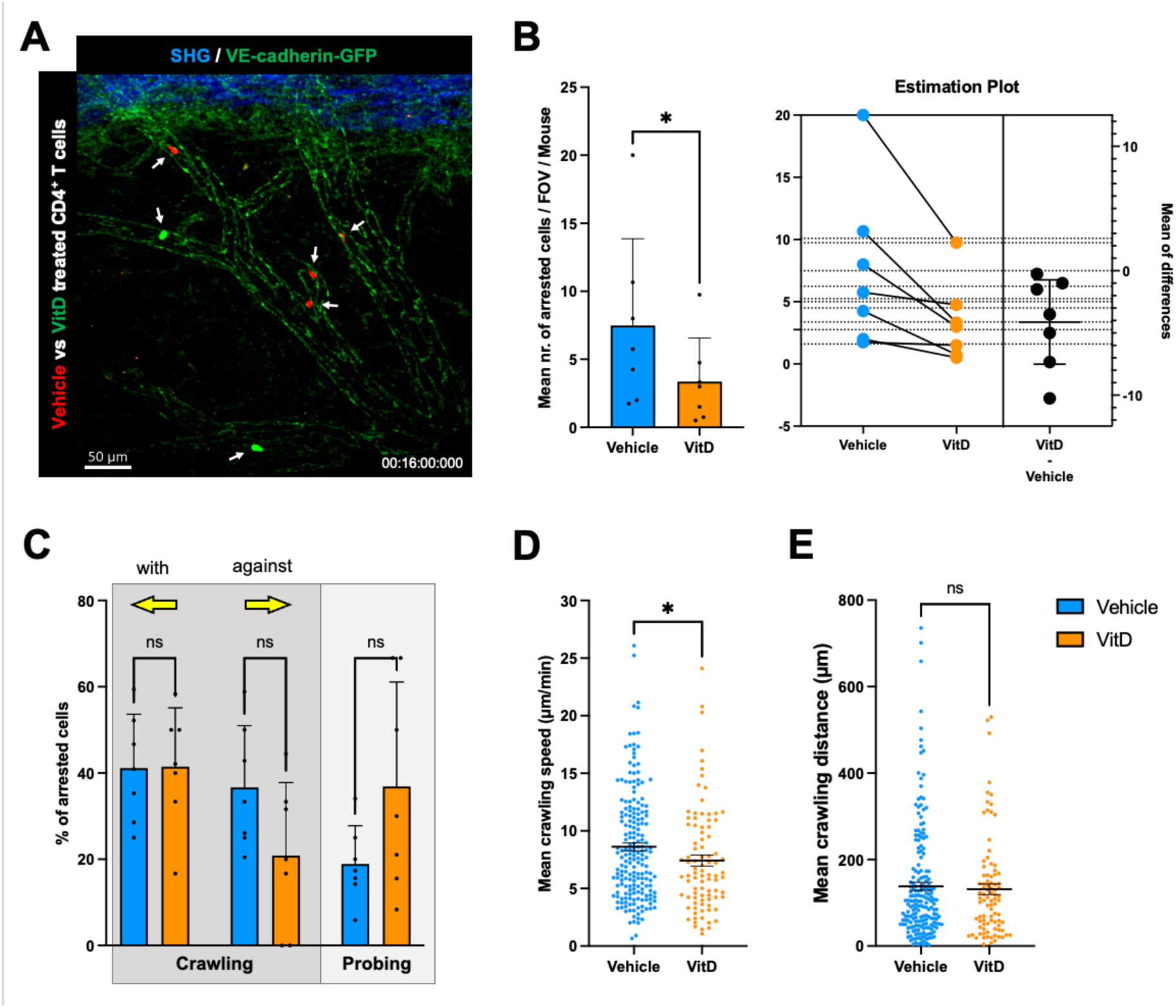
Vitamin D reduces CD4^+^ T-cell arrest to the inflamed BBB *in vivo*. **(A)** Representative picture of human CD4^+^ T cells interacting with cervical spinal cord venules of VE-cadherin-GFP (green) knock-in C57BL/6J mice suffering from active EAE at time point 16 min. 100nM 1,25(OH)2D3 or vehicle control treated CD4^+^ T cells were respectively labelled with CMFDA cell tracker green (green) or cell tracker deep red (red). Autofluorescence resulting from the second harmonic generation (SHG) of the dura matter is depicted in blue. White arrows indicate arrested CD4^+^ T cells on the mouse cervical spinal cord venules. Scale bar = 50 μm. **(B)** Mean number of arrested CD4^+^ T cells treated with 100nM of 1,25(OH)2D3 (orange) or vehicle control (blue) to cervical spinal cord venules of VE-cadherin-GFP knock-in C57BL/6J mice (n = 7) suffering from active EAE (14-17 d post-immunization). **(C)** Quantification of the post-arrest dynamic behavior of CD4^+^ T cells on cervical spinal cord venules during 1h of recording. Behavioral categories were subdivided in crawling and probing, and are shown as fraction of all shear resistant arrested CD4^+^ T cells (100%). Within the category of crawling, we additionally distinguished directionality of the T-cell movement: with or against the flow, depicted with a yellow arrow. **(B, C)** Bar graphs show mean ± SD for 7 movies (mice) of 3 independent EAE experiments. Mean crawling speed **(D)** and distance **(E)** of CD4^+^ T cells on cervical spinal cord venules. **(D, E)** Graph shows mean ± SEM of all arrested CD4^+^ T cells (196 for VitD tretment and 90 for Vehicle control). Each data point represents the velocity and distance of one cell. Statistical analysis: **(B, C)** paired t-test and **(D, E)** Mann-Whitney test (p < 0.05 = *, p < 0.01 = **, p < 0.001 = ***, p < 0.0001 = ****).

### Vitamin D decreases the cell surface expression of α4β1- and αLβ2-integrins on CSF effector CD4 T cells of MS patients

We had previously shown that VitD reduces α4β1-integrin expression on circulating CD4 T cells isolated from PwMS, suggesting that VitD reduces pathogenic CD4 T cell entry into the CNS also in MS.^28^ Hence, we isolated CD4 T cells from the CSF of 5 PwMS and assessed the impact of VitD treatment on expression of the adhesion molecules previously assessed in healthy donor T cells **(Supplementary Fig. 2; Supplementary Table 2)**. Similarly to healthy donor T cells, CSF derived Teff from PwMS showed a homogenous cell surface expression of α4-, α5-, α6-, αL-, β2-integrins and PSGL-1, while two distinct sub-populations with high and low or low/negative cell surface expression were observed for β1- and β7-integrins **(Figure 8A-H).** VitD significantly reduced α4-, αL-, and β2-integrins on all CSF Th subsets compared to vehicle control **(Figure 8A-C)**. In contrast, VitD significantly increased β1-integrins on Th1 cells but reduced it on Th1* and Th17 cells **(Figure 8D)**. Similarly to Th cells from healthy donors, VitD strongly diminished the surface expression of β7-integrins on all CD4 Th subsets from the CSF of PwMS **(Figure 8G)**. α5-integrins were significantly reduced only on Th2 and Th17 cells upon VitD treatment **(Figure 8E)**. Furthermore, no significant impact of VitD on α6-integrins and PSGL-1 was observed **(Figure 8F, H).** Taken together, our results demonstrate that VitD decreases the cell surface expression of important adhesion molecules involved in T cell trafficking in the CNS on pathogenic Teff isolated from the CSF of PwMS. Our data therefore suggest that VitD supplementation may specifically reduce the migration of Teff in the CNS in PwMS.

**Figure 8.**
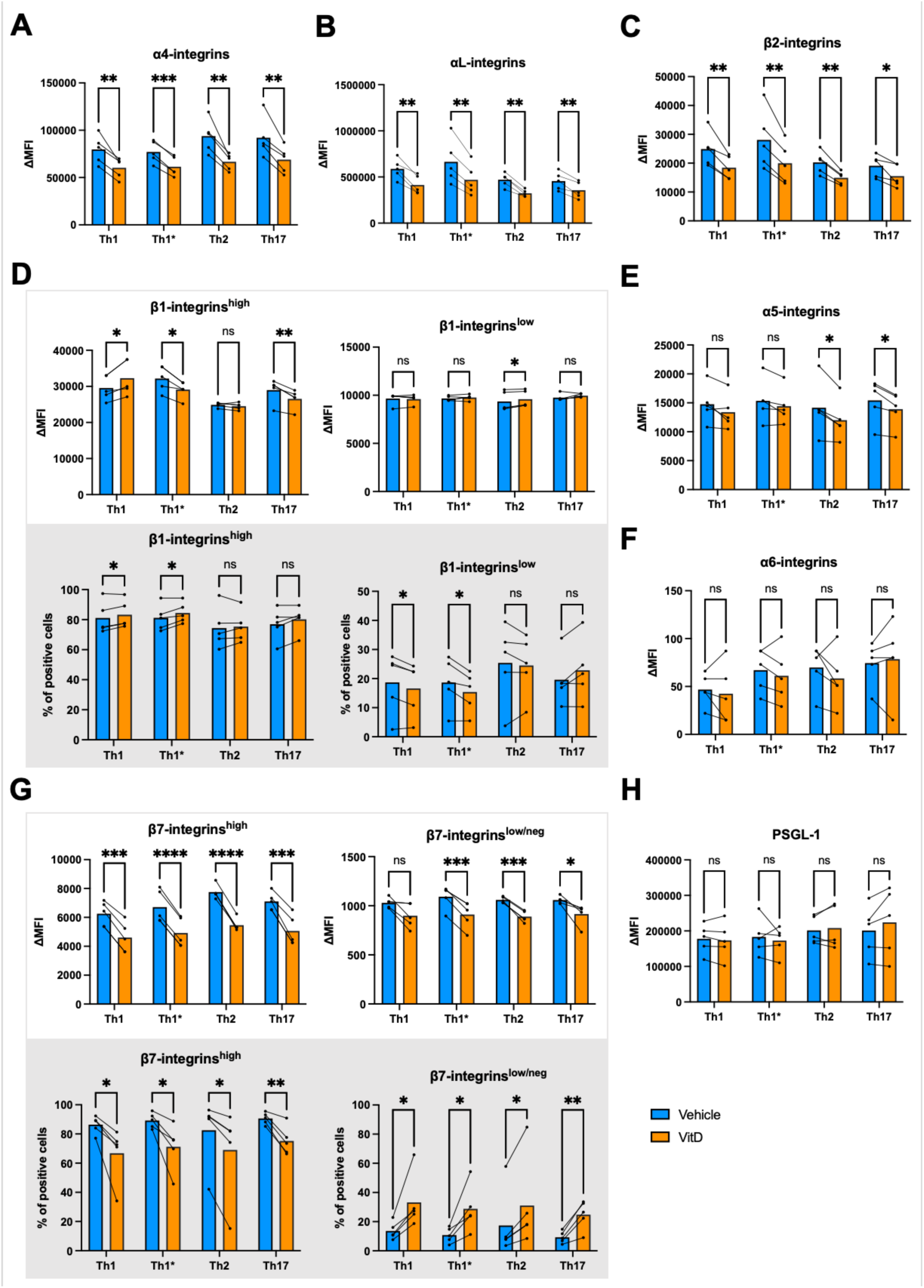
Vitamin D reduces the cell-surface expression of integrins involved in CNS trafficking on pathogenic CD4^+^ T-cell subsets from MS patients. (A-H) Geometric ΔMFI (MFI specific staining - MFI isotype) and percentage of positive cells for the cell-surface expression of different adhesion molecules on different CD4^+^ T-cell subsets (Th1, Th1*, Th2, Th17) from 5 MS patients treated with 100nM of 1,25(OH)2D3 (orange) or vehicle control (blue). Bar graphs show the mean ± SD of 5 independent experiments for the cell-surface expression analysis of α4-integrins **(A)**, αL-integrins **(B)**, β2-integrins **(C)**, β1-integrins **(D)**, α5-integrins **(E)**, α6-integrins **(F)**, β7-integrins **(G**), and PSGL-1 **(H)**. **(D, G)** When two distinct population were identified, CD4^+^ T-cell subsets were subdivided in cells expressing high and low or low/neg cell-surface levels of the different adhesion molecules for analysis. Statistical analysis: paired t-test (p < 0.05 = *, p < 0.01 = **, p < 0.001 = ***, p < 0.0001 = ****).

## Discussion

In the present study, we demonstrate that VitD plays an important role in regulating human T cell migration into the CNS. VitD reduces cell-surface expression of α4β1- and αLβ2-integrins on human CD4 T cells, as previously reported^28,46^, leading to a reduced T-cell arrest to VCAM-1, ICAM-1, and to the inflamed BBB endothelium under physiological flow *in vitro*, as well as *in vivo*. By dissecting the precise mechanisms involved, we demonstrate that VitD specifically impacted on the post-arrest behaviour of T cells on ICAM-1 and on the BBB *in vitro* and *in vivo* by significantly reducing T-cell ability to maintain shear resistant adhesion, by reducing T-cell crawling speed and their ability to crawl against the direction of the flow.

We deciphered the effect of VitD on the multistep migration of T cells across the BBB, showing that VitD decreases the crawling speed of CD4 T cells on the inflamed BBB endothelium *in vitro* and *in vivo*, suggesting also a partially impaired interaction of T cells with the BBB. The interaction of αLβ2-integrin with ICAM-1 is not only important for T-cell crawling but has also been shown to play a role in the process of T-cell diapedesis across the BBB.^47^ As we previously demonstrated, functional blocking of ICAM-1 reduces the migration of T cells across BLEC monolayers *in vitro.*^48^ Despite lower αLβ2-integrin cell-surface levels on VitD-treated CD4 T cells and their reduced interaction with ICAM-1 under physiological flow *in vitro*, the percentage of VitD-treated shear resistant CD4 T cells transmigrating across BLEC monolayers *in vitro* was not significantly reduced, suggesting that other molecular mechanisms might compensate the role of αLβ2-integrins-ICAM-1 interactions in T-cell diapedesis across the BBB endothelium. Unfortunately, we could only observe few T-cell diapedesis events *in vivo* and were therefore unable to make any direct comparison with our *in vitro* observations.

It was previously observed that VitD reduces CCR6 expression, known to regulate the access of murine Th17 cells into the CNS^49^ on human CD4 T cells and suppresses their chemoattraction towards CCL20 *in vitro.*^50^ Whether this can be translated in the context of neuroinflammation *in vivo* remains to be investigated. Furthermore, VitD was shown to reduce the cell-surface expression of CXCR3 on CD4 T cells in EAE, associated with the decreased migration of encephalitogenic T cells into the CNS.^26^ Whether the reduction in CXCR3 cell-surface expression on CD4 T cells in EAE mice is due to a direct or indirect effect of VitD on these cells remains to be investigated. VitD may also affects the migration of recently described pathogenic T-bet+ CXCR3+ B and T cells into the CNS ^51^ ^52^.

The humanized α4-integrin blocking antibody natalizumab (NTZ) is an effective treatment for RRMS^53,54^, suggesting that reduced cell-surface expression of α4-integrins on T cells upon VitD supplementation in RRMS may also lead to similar beneficial effects. VitD supplementation was recently shown to reduce the annual relapse rate in NTZ-treated RRMS patients^55^, suggesting that the combination of NTZ treatment and VitD supplementation may result in better clinical outcomes.

Most importantly, we show that VitD exerts a differential impact on the migration of Teff and Treg across the BBB. VitD significantly reduced the cell-surface levels of α4-integrins, β2-integrins, PSGL-1, and CD99 on the different Teff investigated but not on Treg. Furthermore, VitD reduced the arrest of Th1 but promoted the diapedesis of Treg across the BBB *in vitro*. These data indicate a specificity of action of VitD in the migration of Th subsets involved in MS, and suggest that VitD supplementation in MS might reduce Teff while promoting Treg infiltration into the CNS and diminishing neuroinflammation. Despite the well-established correlation between VitD levels and MS incidence and severity, clinical VitD supplementation trials had mixed outcomes (reviewed in ^12^). However, the recent large D-lay MS trial strongly supports the beneficial effect of VitD supplementation in early MS.^11^

We could not observe any significant effect of VitD on the integrity of the BBB *in vitro*, indicating that in our model, VitD directly modulates T cells and impacts their arrest and interaction with the BBB. This is in contrast with a previous study that reported changes in TEER upon VitD treatment of human brain microvascular endothelial cells (HBMECs) incubated with TNFα, another proinflammatory molecule, or serum from MS patients *in vitro.*^56^ BLECs may be a more relevant *in vitro* model of the BBB than HBMECs by responding to co-culture with pericytes or pericyte-conditioned medium with improved barrier properties such as higher TEER and lower permeability to small molecular tracers and expression of BBB specific molecules alone.^32^ Other studies showed that VitD treatment reduces the leakage of different tracers into the CNS parenchyma during EAE.^26,57^ Similarly, fewer new Gd enhancing lesions were observed in MS patients supplemented with cholecalciferol compared to placebo group.^58–60^ Whether these observations are due to a direct or indirect effect of VitD on the BBB endothelium remains to be elucidated.

Overall, we show that VitD regulates T cell migration into the CNS and exerts direct effects on T cells by impeding Teff migration while allowing Treg migration into the CNS. These data, combined with the knowledge of the immunoregulatory role of VitD and the positive effect of VitD treatment on EAE, further support a key role for VitD in reducing MS disease activity and progression.

## Data availability

The original data used and/or analyzed during the current study are available from the corresponding authors upon request.

## Acknowledgements

We are very grateful to the patients who volunteered for this study. We thank Fatima L’faqihi and Anne Laure Iscache (INFINITY, Toulouse) for their help with the flow cytometry, and the Immunomonitoring platform at Infinity. The authors are thankful for helpful discussion with Drs. Frederick Masson, Abdel Saoudi, Roland Liblau. The authors have no conflict of interest.

## Funding

This research has been supported by the Swiss National Science Foundation (SNSF) grant 310030E_189312 to BE, the Swiss MS Society to BE and the SNFS grant CRSII3_154483 to BE, FS and RM, and by European Research Council Advanced grant ERC-2013-ADG 340733 to R.M. Further funding was provided by the Agence Nationale de la Recherche (ANR) grant ANR-19-CE14-0043, France SEP, the UK Multiple Sclerosis Society (859/07 and MS 41) to ALA. Additional funding was provided by the Eugène Devic EDMUS Foundation against multiple sclerosis in partnership with ARSEP foundation grant to ALA and BE, and the Hubert Curien/Germaine de Stael scheme to BE/ALA, by the Bangerter Rhyner Foundation to BE and the Swiss MS Society to AM. JK obtained a travel grant from the Edinburgh Neuroscience funds. HN has been supported by the Uehara Memorial Foundation and an ECTRIMS fellowship. NS, PO, and HW were partly supported by grants from the Deutsche Forschungsgemeinschaft (DFG, SFB128 A10 and B01) and the IZKF Muenster (Project Wie3/009/16).

## Competing interests

The authors report no competing interests.

## Supplementary material

Supplementary material is available at *Brain* online.

## Notes

### Competing Interest Statement

The authors have declared no competing interest.

