## Supplemental data for "Vitamin D differentially modulates effector and regulatory T-cell migration across the blood-brain barrier"

**Affiliation:**

**Present address:**

### Supplementary Figures

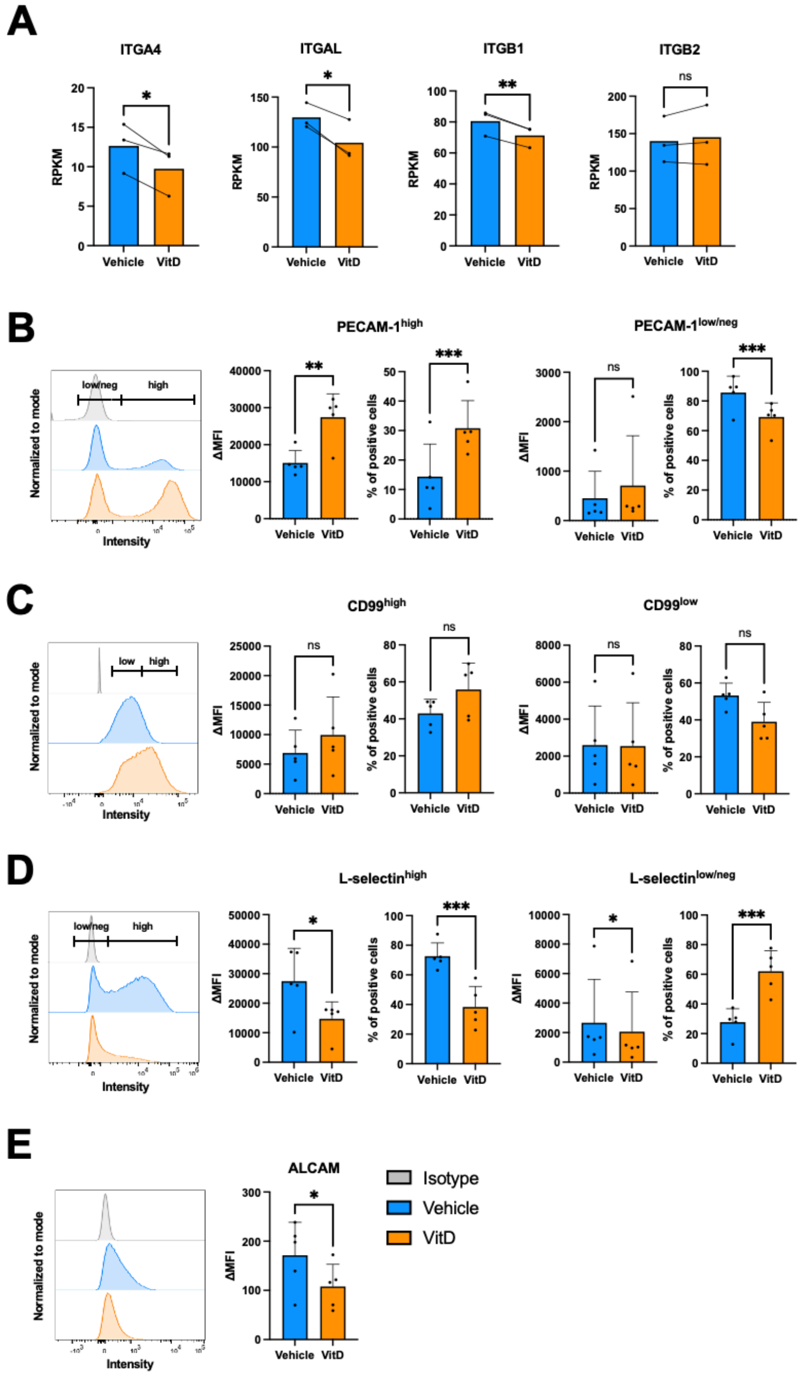


**Supplementary Figure 1. Vitamin D regulates the cell-surface expression of other important adhesion molecules for $CD4^+$ T-cell extravasation. (A)** Normalized count for RNA expression of ITGA4, ITGAL, ITGB1 and ITGB2 from previously published RNA-seq data of CD3/CD28 activated human CD4 T cells in the presence or absence of VitD (GEO GSE154741).[1] **(B-E)** Representative histogram plots, geometric ΔMFI (MFI specific staining - MFI isotype) and percentage of positive cells for the cell-surface expression of different adhesion molecules on CD4 T cells treated with 100nM of $1,25(OH)_2D3$ (orange) or vehicle control (blue). Isotype control for each flow cytometry staining is shown in grey in each histogram plot. Bar graphs show the mean ± SD of 5 independent experiments (healthy donors) for the cell-surface expression analysis of PECAM-1 **(B)**, CD99 **(C)**, L-selectin **(D)**, and ALCAM **(E). (B-D)** When two distinct populations were identified, CD4 T cells were subdivided for the analysis in cells expressing high and low or low/neg cell-surface levels of the respective adhesion molecules. The gating strategy for high and low or low/neg populations is displayed in each respective histogram plot. Statistical analysis: paired t-test ($p < 0.05$ = *, $p < 0.01$ = **, $p < 0.001$ = ***, $p < 0.0001$ = ****).

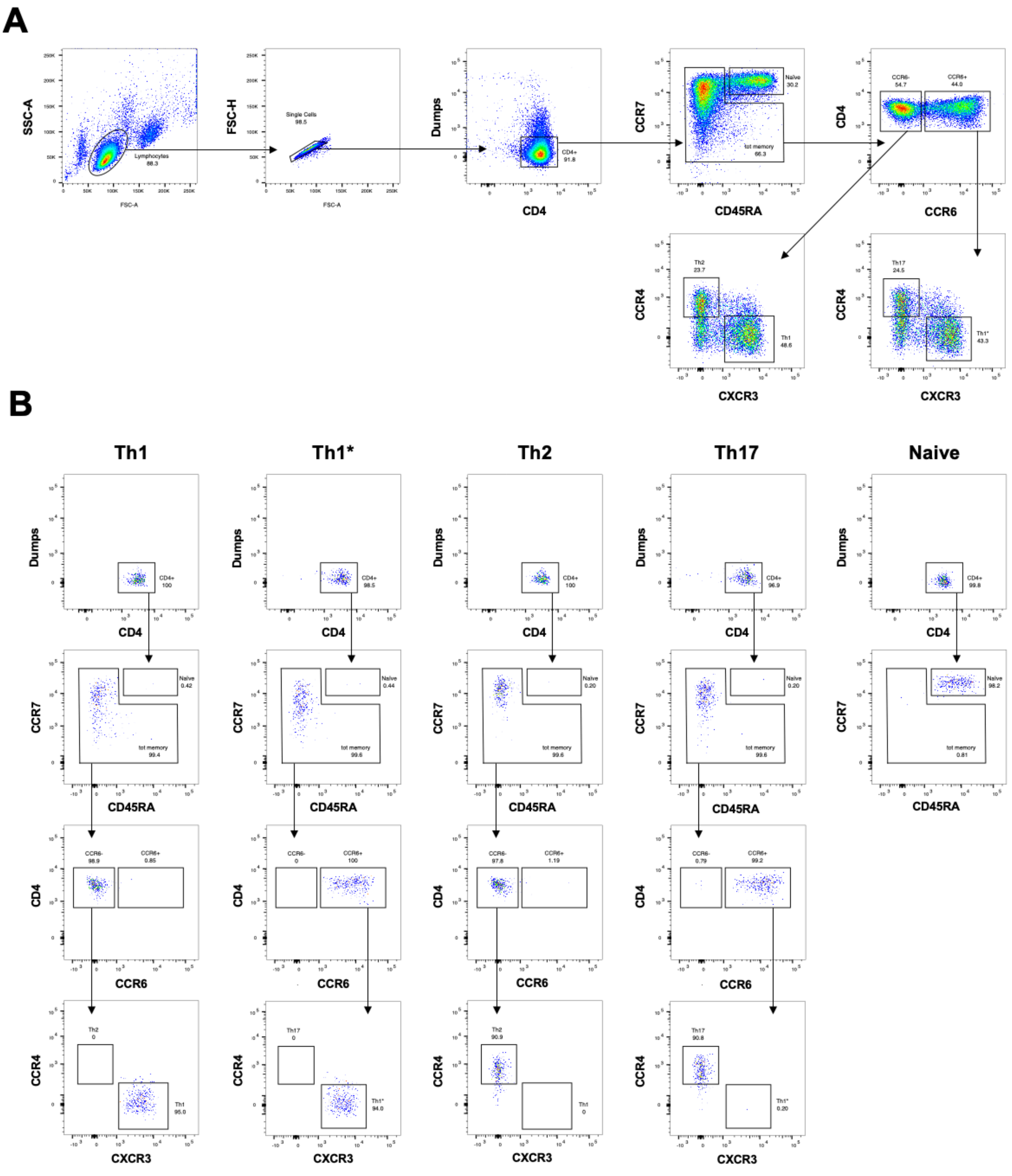


**Supplementary Figure 2. Isolation and sorting of different human effector/memory CD4$^{+}$ Th-subsets. (A)** Representative gating strategy for the fluorescence activated cell-sorting (FACS) of human effector/memory CD4$^{+}$ Th1, Th1*, Th2, and Th17 cells. **(B)** Th-cell purity after FACS.

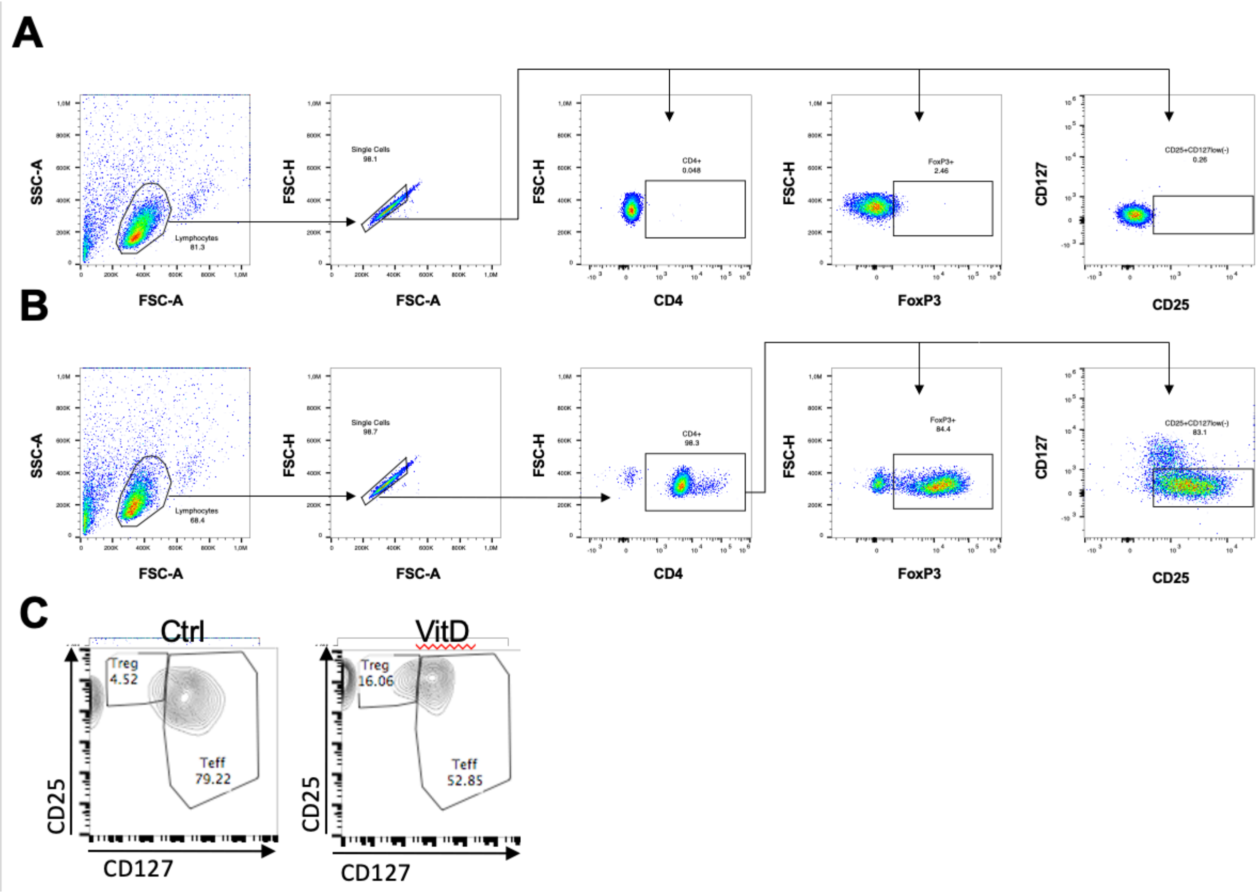


**Supplementary Figure 3. CD4$^+$FoxP3$^+$ regulatory T-cell purity after isolation.** Flow cytometry analysis for isotype control **(A)**, isolated CD4$^+$CD25$^+$CD127$^{low/-}$ T cells **(B)** after magnetic cell-sorting. **(C)** Representative density plot with the percentage of transmigrated effector (CD25$^+$CD127$^+$) and regulatory (CD25$^+$CD127$^{low}$) CD4$^+$ T cells after vehicle (Ctrl) and 1,25(OH)$_2$D3 treatment.

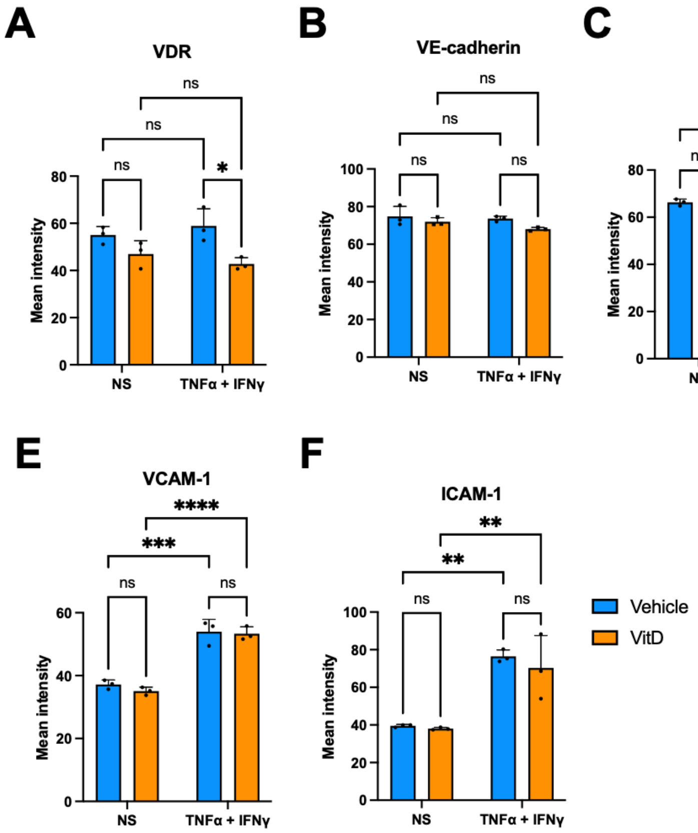


**Supplementary Figure 4. Vitamin D does not change the integrity and cell-surface adhesion molecule expression of VCAM-1 and ICAM-1 of the BBB *in vitro*.** Mean intensity of the florescent signal from the staining for VDR **(A)**, VE-cadherin **(B)**, ZO-1 **(C)**, claudin-5 **(D)**, VCAM-1 **(E)**, and ICAM-1 **(F)** of NS and stimulated BLECs pre-treated with 100nM of 1,25$(OH)_2D3$ (orange) or vehicle control (blue). Bar graphs show mean ± SD of 3 independent experiments. Statistical analysis: **(B-G)** Two-way ANOVA followed by Šídák's multiple comparisons test ($p < 0.05$ = *, $p < 0.01$ = **, $p < 0.001$ = ***, $p < 0.0001$ = ****).

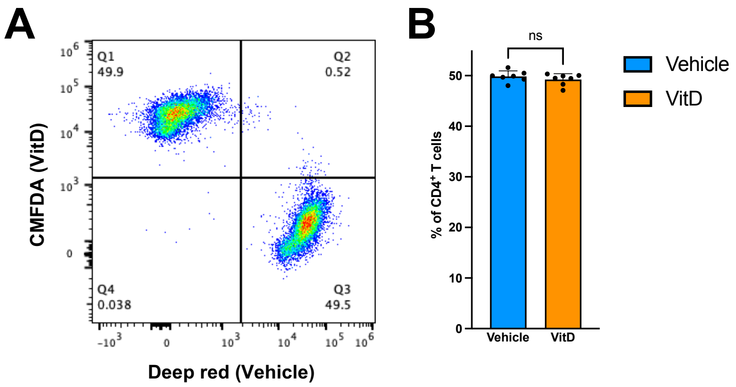


**Supplementary Figure 5. Proportion of vitamin D and vehicle treated human CD4$^+$ T cells injected in mice for 2-photon intravital microscopy. (A)** Representative flow cytometry dot plot (left) and **(B)** percentage of 1,25(OH)$_2$D3 (blue) or vehicle control (orange) treated CD4$^+$ T cells (right) injected systemically via carotid artery catheter. Bar graphs show mean ± SD of 7 experiments**.** Statistical analysis: unpaired t-test ($p < 0.05$ = *, $p < 0.01$ = **, $p < 0.001$ = ***, $p < 0.0001$ = ****).

**Supplementary Movies**

**Supplementary Movie 1.**

Representative zoomed-in time lapse videos of CMFDA (Cell Tracker Green) prelabelled human $CD4^+$ T-cell interaction with VCAM-1 (1.54 μg/mL) under physiological flow. Vehicle control, 1,25$(OH)_2$D3 treatment, and α4-integrins blocking conditions are shown respectively from the left to the right. 10 min of recording are shown: 4 min of accumulation phase (0.1 dyne/$cm^2$) and 6 min of physiological shear stress (1.5 dyne/$cm^2$). The overlay of phase contrast and green-fluorescent channels is shown. Time is indicated as minutes:seconds. Flow direction is illustrated by an arrow (yellow). Scale bar = 50 μm.

**Supplementary Movie 2.**

Representative zoomed-in time lapse videos of CMFDA (Cell Tracker Green) prelabelled human $CD4^+$ T-cell interaction with ICAM-1 (1.14 μg/mL) under physiological flow. Vehicle control, 1,25$(OH)_2$D3 treatment, and β2-integrins blocking conditions are shown respectively from the left to the right. 10 min of recording are shown: 4 min of accumulation phase (0.1 dyne/$cm^2$) and 6 min of physiological shear stress (1.5 dyne/$cm^2$). The overlay of phase contrast and green-fluorescent channels is shown. Time is indicated as minutes:seconds. Flow direction is illustrated by an arrow (yellow). Scale bar = 50 μm.

**Supplementary Movie 3.**

Representative time lapse videos of the x/y diagrams of human $CD4^+$ T-cell crawling tracks on equimolar concentrations of immobilized recombinant ICAM-1 (1.14 μg/mL) **(A)** and VCAM-1 (1.54 μg/mL) **(B)** under physiological flow. Vehicle control and 1,25$(OH)_2$D3 treatment conditions are shown on the left and on the right respectively. Crawling tracks over 5 min and 30 s of physiological shear stress (1.5 dyne/$cm^2$) are displayed. For each track, the site of arrest was set to the center point of the respective diagram and is shown by the intersection of two blue lines. End points of tracks are indicated by a red dot. Time is indicated as minutes:seconds. Flow direction is illustrated by an arrow (yellow).

**Supplementary Movie 4.**

Representative zoomed-in time lapse videos of CMFDA prelabelled (Cell Tracker Green) human $CD4^+$ T-cell interaction with TNFα/IFNγ stimulated BLECs under physiological flow. 1,25$(OH)_2$D3, vehicle control, and α4- and -β2-integrins blocking conditions are shown respectively from the left to the right. 24 min of recording are shown: 4 min of accumulation phase (0.1 dyne/$cm^2$) and 20 min of physiological shear stress (1.5 dyne/cm$^2$). The overlay of phase contrast and green-fluorescent channels is shown. Time is indicated as minutes:seconds. Flow direction is illustrated by an arrow (yellow). Scale bar = 50 µm.

**Supplementary Movie 5.**

Representative time lapse videos of the x/y diagrams of human CD4$^+$ T-cell crawling tracks on BLECs under physiological flow. Vehicle control and 1,25(OH)$_2$D3 treatment conditions are shown on the left and the right respectively. Crawling tracks over 20 min of physiological shear stress (1.5 dyne/cm$^2$) are displayed. For each track, the site of arrest was set to the center point of the respective diagram and is shown by the intersection of two blue lines. End points of tracks are indicated by a red dot. Time is indicated as minutes:seconds. Flow direction is illustrated by an arrow (yellow).

**Supplementary Movie 6.**

Representative time lapse videos of human CD4$^+$ T-cell interaction with the cervical spinal cord venules of a VE-cadherin-GFP reporter mice suffering active EAE (d15 post immunization). 1,25(OH)$_2$D3 treated CD4$^+$ T cells were prelabelled CMFDA cell tracker green, while vehicle treated were prelabelled with cell tracker deep red and are shown in green and red respectively. VE-cadherin-GFP and the autofluorescence of the second harmonic generation (SHG) are displayed in green and blue respectively. 1 hour of recording is shown. The video shows the maximum intensity projection of the blue, green and far-red-fluorescent channels overlayed. Time is indicated as hours:minutes:seconds:milliseconds. Scale bar = 100 µm.

### Supplementary Tables

#### Supplementary Table 1. Characteristics of CSF samples from MS patients

| donor | type | gender | age | disease duration (month) | treatment before CSF | CSF (cells/μL) | Total Protein CSF (mg/L) | IgG index | oligoclonal band | clinical disease activity | MRI Gd enhancement |
|---|---|---|---|---|---|---|---|---|---|---|---|
| 1 | CIS | F | 19 | 0 | - | 2 | 233 | 0.50 | + | + | - |
| 2 | RRMS | F | 25 | 10 | - | 6 | 470 | 0.64 | + | - | - |
| 3 | RRMS | F | 33 | 2 | - | 7 | 338 | 1.96 | + | + | + |
| 4 | RRMS | M | 50 | 34 | - | 3 | 486 | 0.98 | + | - | + |
| 5 | RRMS | M | 22 | 13 | - | 7 | 263 | 0.52 | + | + | + |

#### Supplementary Table 2. Antibody list for flow cytometry analysis of CD4 and CD8 T-cell purity

| Target | Fluorophore | Clone | Provider | Cat. Number | Isotype |
|---|---|---|---|---|---|
| **anti-human CD3** | AF700 | OKT3 | Biolegend | 317340 | m-IgG2a, κ |
| **anti-human CD4** | FITC | RPA-T430 | Biolegend | 305006 | m-IgG1, κ |
| **anti-human CD3** | APC | HIT3a | Biolegend | 300312 | m-IgG2a, κ |
| **anti-human CD8** | APC-Cy7 | SK1 | Biolegend | 344714 | m-IgG1, κ |

| Isotype controls | Fluorophore | Clone | Provider | Cat. Number |
|---|---|---|---|---|
| **m-IgG2a, κ** | AF700 | MOPC-173 | Biolegend | 400247 |
| **m-IgG1, κ** | FITC | MOPC-21 | Biolegend | 400108 |
| **m-IgG2a, κ** | APC | MOPC-173 | Biolegend | 400420 |
| **m-IgG1, κ** | APC-Cy7 | MOPC-21 | BD Biosciences | 557873 |

#### Supplementary Table 3. Antibody list for fluorescence-activated cell sorting and purity of different CD4 Th subsets

| Target | Fluorophore | Clone | Provider | Cat. Number | Isotype |
|---|---|---|---|---|---|
| **anti-human CD4** | PE-Tr | S3.5 | ThermoFischer Scientific | MHCD0417 | m-IgG2a, κ |
| **anti-human CD45RA** | Qd655 | MEM-56 | ThermoFischer Scientific | Q10069 | m-IgG2b, κ |
| **anti-human CD183 (CXCR3)** | AF647 | G025H7 | Biolegend | 353711 | m-IgG1, κ |
| **anti-human CD194 (CCR4)** | PE-Cy7 | 1G1 | BD Biosciences | 557864 | m-IgG1, κ |
| **anti-human CD196 (CCR6)** | PE | 11A9 | BD Biosciences | 559562 | m-IgG1, κ |
| **anti-human CD197 (CCR7)** | BV421 | G043H7 | Biolegend | 353208 | m-IgG2a, κ |
| **anti-human CD8** | PE-Cy5 | B9.11 | Beckman Coulter | A07758 | m-IgG1, κ |
| **anti-human CD14** | PE-Cy5 | RMO52 | Beckman Coulter | A07765 | m-IgG2a, κ |
| **anti-human CD19** | PE-Cy5 | HIB19 | ThermoFischer Scientific | 15-0199-42 | m-IgG1, κ |
| **anti-human CD25** | PE-Cy5 | B1.49.9 | Beckman Coulter | IM2646 | m-IgG2a, κ |
| **anti-human CD56** | PE-Cy5 | N901 | Beckman Coulter | A07789 | m-IgG1, κ |

**Supplementary Table 4. Antibody list for flow cytometry analysis of CD4 Treg purity**

| Target | Fluorophore | Clone | Provider | Cat. Number | Isotype |
|---|---|---|---|---|---|
| **anti-human CD3** | AF700 | OKT3 | Biolegend | 317340 | m-IgG2a, *κ* |
| **anti-human CD4** | FITC | RPA-T430 | Biolegend | 305006 | m-IgG1, *κ* |
| **anti-human CD25** | BV605 | BC96 | Biolegend | 302632 | m-IgG1, *κ* |
| **anti-human CD127** | BV421 | HIL-7R-M21 | BD Biosciences | 562436 | m-IgG1, *κ* |
| **anti-human FoxP3** | PE | 206D | Biolegend | 320108 | m-IgG1, *κ* |

| Isotype controls | Fluorophore | Clone | Provider | Cat. Number |
|---|---|---|---|---|
| **m-IgG2a, *κ*** | AF700 | MOPC-173 | Biolegend | 400247 |
| **m-IgG1, *κ*** | FITC | MOPC-21 | Biolegend | 400108 |
| **m-IgG1, *κ*** | BV605 | MOPC-21 | Biolegend | 400161 |
| **m-IgG1, *κ*** | BV421 | X40 | BD Biosciences | 562438 |
| **m-IgG1, *κ*** | PE | MOPC-21 | Biolegend | 400112 |

**Supplementary Table 5. Antibody list for flow cytometry analysis of adhesion molecules surface expression on BLECs**

| Target | Fluorophore | Clone | Provider | Cat. Number | Isotype |
|---|---|---|---|---|---|
| **anti-human CD54 (ICAM-1)** | BV421 | HA58 | BD Biosciences | 564077 | m-IgG1, *κ* |
| **anti-human CD106 (VCAM-1)** | FITC | 51-10C9 | BD Biosciences | 551146 | m-IgG1, *κ* |
| **anti-human CD144 (VE-cadherin)** | PerCP-Cy5.5 | 55-7H1 | BD Biosciences | 561566 | m-IgG1, *κ* |
| **anti-human CD102 (ICAM-2)** | PE | CBR-IC2/2 | BD Biosciences | 558080 | m-IgG2a, *κ* |
| **anti-human CD99** | PE-Cy7 | 3B2/TA8 | Biolegend | 371314 | m-IgG2a, *κ* |
| **anti-human CD31 (PECAM-1)** | APC-Cy7 | WM59 | BD Biosciences | 563653 | m-IgG1, *κ* |

| Isotype controls | Fluorophore | Clone | Provider | Cat. Number |
|---|---|---|---|---|
| **m-IgG1, *κ*** | BV421 | X40 | BD Biosciences | 562438 |
| **m-IgG1, *κ*** | FITC | MOPC-21 | Biolegend | 400108 |
| **m-IgG1, *κ*** | PerCP-Cy5.5 | MOPC-21 | Biolegend | 400150 |
| **m-IgG1, *κ*** | APC-Cy7 | MOPC-21 | BD Biosciences | 557873 |
| **m-IgG2a, *κ*** | PE | eBM2a | Invitrogen | 12-4724-42 |
| **m-IgG2a, *κ*** | PE-Cy7 | MOPC-173 | Biolegend | 400232 |

**Supplementary Table 6. Antibody list for flow cytometry analysis of adhesion molecules expression on human T cells**

| Target | Fluorophore | Clone | Provider | Cat. Number | Isotype |
|---|---|---|---|---|---|
| **anti-human CD29 (β1-integrin)** | APC | MAR4 | BD Biosciences | 559883 | m-IgG1, κ |
| **anti-human CD49d (α4-integrin)** | APC-Cy7 | 9F10 | Biolegend | 304328 | m-IgG1, κ |
| **anti-human β7-integrin** | BV650 | FIB504 | BD Biosciences | 564284 | rt-IgG2a, κ |
| **anti-human CD18 (β2-integrin)** | FITC | 6.7 | BD Biosciences | 555923 | m-IgG1, κ |
| **anti-human CD103 (αE-integrin)** | BV711 | Ber-ACT8 | BD Biosciences | 563162 | m-IgG1, κ |
| **anti-human CD62L (L-selectin)** | APC | DREG-56 | BD Biosciences | 559772 | m-IgG1, κ |
| **anti-human PSGL-1 (CD162)** | PE | KPL-1 | Biolegend | 328806 | m-IgG1, κ |
| **anti-human CD11a (αL-integrin)** | Pe-Cy7 | HI111 | BD Biosciences | 561387 | m-IgG1, κ |
| **anti-human CD51 (αV-integrin)** | FITC | NKI-M9 | Biolegend | 327908 | m-IgG2a, κ |
| **anti-human CD31 (PECAM-1)** | APC-Cy7 | WM59 | BD Biosciences | 563653 | m-IgG1, κ |
| **anti-human CD99** | FITC | HEC2 | Biolegend | 398208 | m-IgG1, κ |
| **anti-human CD166 (ALCAM)** | PerCP-Cy5.5 | 3A6 | Biolegend | 343908 | m-IgG1, κ |
| **anti-human CD49e (α5-integrin)** | APC | MKI-SAM1 | Biolegend | 328012 | m-IgG2b, κ |
| **anti-human CD49f (α6-integrin)** | FITC | GoH3 | Biolegend | 313606 | rat-IgG2a, κ |

| Isotype controls | Fluorophore | Clone | Provider | Cat. Number |
|---|---|---|---|---|
| **m-IgG1, κ** | APC | MOPC-21 | BD Biosciences | 555751 |
| **m-IgG1, κ** | APC-Cy7 | MOPC-21 | BD Biosciences | 557873 |
| **m-IgG1, κ** | PE | MOPC-21 | Biolegend | 400112 |
| **m-IgG1, κ** | PerCP-Cy5.5 | MOPC-21 | Biolegend | 400150 |
| **m-IgG1, κ** | FITC | MOPC-21 | Biolegend | 400108 |
| **m-IgG1, κ** | PE-Cy7 | MOPC-21 | BD Biosciences | 557872 |
| **m-IgG1, κ** | BV421 | X40 | BD Biosciences | 562438 |
| **m-IgG1, κ** | BV711 | MOPC-21 | Biolegend | 40016 |
| **m-IgG2a, κ** | FITC | MOPC-173 | Biolegend | 981902 |
| **m-IgG2b, κ** | APC | MPC-11 | Biolegend | 400322 |
| **rt-IgG2a, κ** | BV650 | R35-95 | BD Biosciences | 563144 |
| **rt-IgG2a, κ** | FITC | RTK2758 | Biolegend | 400506 |

**Supplementary Table 7. Antibody list for immunofluorescence staining of adhesion and junctional molecules of BLECs**

| Target | Clone | Provider | Cat. Number | Isotype | Secondary Antibody |
|---|---|---|---|---|---|
| **anti-human CD54 (ICAM-1)** | HA58 | Biolegend | 353102 | m-IgG1, κ | AF488 Donkey Anti-Mouse Affine Pure IgG (H+L) |
| **anti-human CD 106 (VCAM-1)** | 51-10C9 | R&D systems | 555645 | m-IgG1, κ | AF488 Donkey Anti-Mouse Affine Pure IgG (H+L) |
| **anti-human ZO-1** | polyclonal | Invitrogen | 40-2200 | rb-IgG | Cy3 AffiniPure Donkey Anti-Rabbit IgG (H+L) |
| **anti-human VE-cadherin** | clone F-8 | Santa Cruz Biotechnology | sc-9989 | m-IgG1, κ | AF488 Donkey Anti-Mouse Affine Pure IgG (H+L) |
| **anti-human claudin-5** | 4C3C2 | Invitrogen | 35-2500 | m-IgG1, κ | Cy3 AffiniPure F(ab')2 Fragment Goat Anti-Mouse IgG |
| **anti-human VDR** | 9A7 | Invitrogen | MA1-710 | rt-IgG2b, κ | Cy3 AffiniPure Donkey Anti-Rat IgG (H+L) |
| **anti-human fibronectin** | polyclonal | DAKO | A0245 | rb-IgG | Cy3 AffiniPure Donkey Anti-Rabbit IgG (H+L) |

**Supplementary Table 8. Antibody list for flow cytometry analysis of transmigrated T cells across BLECs**

| Target | Fluorophore | Clone | Provider | Cat. Number | Isotype |
|---|---|---|---|---|---|
| **anti-human CD25** | PE | BC96 | Biolegend | 302605 | m-IgG1, κ |
| **anti-human CD127** | APC | A019P5 | Biolegend | | m-IgG1, κ |

1 Chauss, D. *et al.* Autocrine vitamin D signaling switches off pro-inflammatory programs of T(H)1 cells. *Nat Immunol* **23**, 62-74 (2022). https://doi.org:10.1038/s41590-021-01080-3
